# Immune Checkpoint Therapy Drives Maturation of a Cellular Neighborhood Nucleated by T Cell-APC Triads Enabling Spatially Compartmentalized Tumor Immunity

**DOI:** 10.64898/2026.04.08.716779

**Authors:** Ruan F. Vieira Medrano, Vladimir Sukhov, Kevin Hoffer-Hawlik, Bar Rozenman, Cora D. Arthur, Fei Han, Bassem Ben Cheikh, Hussein Sultan, Yoshiko Takeuchi, Samuel Ameh, Derek J. Theisen, Yuang Song, Heather N. Kohlmiller, J. Michael White, Alexey Sergushichev, Bernd H. Zinselmeyer, Patrick Leinert, Wm. Patrick Leinert, Kathleen C.F. Sheehan, Garry Nolan, Elham Azizi, Maxim N. Artyomov, Robert D. Schreiber

## Abstract

Spatially organized immune hubs of T cells and antigen-presenting cells (APCs) have been linked to immune checkpoint therapy (ICT) efficacy, yet the mechanisms underlying their function remain unclear. Using CODEX multiplex imaging, we longitudinally characterized the dynamic evolution of intratumoral cellular neighborhoods (CN) defined by “triad” interactions of CD4⁺ and CD8⁺ T cells with two distinct myeloid APC populations: cDC1s and IFNγ–activated macrophages. We termed this CN the immunity-promoting CN (IP-CN) and tracked its progressive development during tumor rejection induced by α-CTLA-4/α-PD-1 therapy. A coordinated IFNγ and TNFα signaling signature accompanied the IP-CN assembly. Over time, the IP-CN underwent functional maturation, forming specialized sub-neighborhoods that compartmentalized proliferating T cells at the tumor periphery versus cytotoxic T effector cells interacting with tumor cell targets. Our findings reveal a spatiotemporal mechanism by which the IP-CN sustains and amplifies cytotoxic T cell responses, demonstrating how T cell–APC neighborhoods orchestrate tumor immunity.

## INTRODUCTION

Advancements in our understanding of the mechanisms employed by the immune system to control tumor outgrowth have led to groundbreaking discoveries in cancer immunotherapy^1, 2, 3, 4, 5^. This progress is particularly underscored by the clinical success of Immune-Checkpoint Cancer Therapy (ICT) targeting the Cytotoxic T-lymphocyte Associated Antigen-4 (CTLA-4) and Programmed Death-1 (PD-1) immune checkpoint pathways, which have revolutionized cancer treatment, sometimes providing durable responses in patients with different types of cancer ^6, 7, 8^. However, a significant fraction of tumor-bearing individuals fail to respond to therapy^9^ and thus, a deeper understanding of the mechanisms underlying successful cancer immunotherapy is critically needed.

Previous work in our lab has shed light on this issue by defining mechanisms of successful tumor rejection^10, 11, 12^. We demonstrated that optimal immune responses to developing and established tumors required that (a) tumor cells express antigens for both CD4^+^ and CD8^+^ T cells; (b) the host was able to induce tumor-specific CD4^+^ and CD8^+^ T cells; and (c) both requirements were met even in tumors that lacked the expression of MHC-II proteins^13^. Furthermore, using CyTOF and scRNAseq analysis of tumor-infiltrating CD45^+^ cells, we showed that successful ICT induced a profound remodeling of the lymphoid compartment, which was marked by CD4^+^ and CD8^+^ T cell activation and proliferative responses^14^. This work also revealed that the myeloid/macrophage compartment also concomitantly underwent major remodeling such that intratumoral CX3CR1^+^ macrophages expanded during tumor progression, but rapidly disappeared following ICT, whereas IFNγ-induced iNOS^+^ macrophages increased during tumor rejection^14^.

Recent studies highlight the critical role of immune cell spatial organization in determining outcomes following immunotherapy^15, 16, 17, 18^. Most notably, tertiary lymphoid structures (TLS) are well-established niches for cellular crosstalk associated with a favorable prognosis^19, 20, 21^. Analogous to TLS, additional forms of immune neighborhoods composed of T cells and antigen-presenting cells (APCs) have also been reported^22, 23^. However, our understanding of how these structures are formed and change over time remains limited. Importantly, most of our current insights derive from spatial analyses performed at single time points or under conditions that result in only partial therapeutic responses. As a result, the spatiotemporal mechanisms governing the functional role of these T cell-APC hubs during successful immunotherapy remain largely undefined.

To address this deficiency, we applied a longitudinal spatial-proteomic approach herein to investigate the changes, over time, that occur in immune cell organization induced by ICT in a fully responsive mouse tumor model. Mice bearing syngeneic orthotopic T3 tumors were treated with control monoclonal antibodies (mAb) or the dual ICT combination of α-CTLA-4 and α-PD-1 (herein referred to as α-CTLA-4/α-PD-1). Tumors were harvested at multiple time points and subjected to CODEX multiplex imaging^24, 25^, allowing us to profile, at single-cell resolution, surface and intracellular proteins in fresh-frozen tissue sections. We found that T3 tumors are organized into seven distinct cellular neighborhoods (CNs) and that α-CTLA-4/α-PD-1 promotes dramatic changes in their trajectories. Specifically, tumor rejection was orchestrated through the assembly, expansion, and maturation of an immunity-promoting CN (IP-CN) composed of CD4⁺ and CD8⁺ T cells forming three-cell clusters (triads) with cDC1s or IFNγ–activated macrophages. We observed that these triads form early in the α-CTLA-4/α-PD-1-induced rejection process and increase in frequency as the IP-CN expands, indicating that continuous T cell/APC engagement is an important feature nucleating the response. We also observed that the IP-CN supports the intratumoral induction of T cell proliferation and effector function. But over time, these functions become spatially compartmentalized into specialized sub-neighborhoods, with proliferating T cells enriched with triad-forming APCs at the tumor tissue periphery as cytotoxic T cells progressively move inward, targeting cancer cells at the tumor boundary CN. Overall, this study provides a spatiotemporal mechanism for the central role that the IP-CN has in organizing tumor immunity. Our findings establish maturation of immune neighborhoods and functional compartmentalization as novel principles underlying successful ICT.

## RESULTS

### Spatial Proteomic Profiling to Monitor Successful Immune Checkpoint Therapy in T3 Tumors

For this study, we utilized our well-characterized T3 MCA sarcoma line, which grows progressively when injected orthotopically into syngeneic wild-type mice but is uniformly rejected with consistent kinetics upon treatment with α-CTLA-4/α-PD-1^10, 13^ in a manner requiring tumor neoantigen-specific CD4^+^ and CD8^+^ T cells (herein recapitulated in **Figure S1A**). We focused our study on the effects induced by α-CTLA-4/α-PD-1 because we had previously established that this treatment induced stronger anti-tumor responses than single-agent therapy^14, 26^ and thus would provide a better model for successful antitumor responses.

To map the spatiotemporal dynamics of the T cell antitumor response, we harvested T3 tumors at days (d) 7, 9, 10, 11 and 13 post-transplant, which were timepoints known to correspond to critical functional changes that we had previously reported^14^. Tumors were fresh-frozen, cut into 8μm sections encompassing the entire tumor-immune interface, and subsequently transferred onto coverslips for staining **(Figure 1A)** with a panel of 32 bar-coded CODEX Abs **(Figure 1B)**. In total, our dataset was derived from 49 tumors (individually displayed in **Figure S2**). Detection of tumor cells was achieved using a mAb specific for CD140a [platelet-derived growth factor receptor-α (PDGF-α)] that was highly expressed by T3 tumor cells **(Figure S1B)**. The expected antigen staining patterns for all antibodies are displayed in **Figures S3A** and **S3B**.

**Figure 1.**
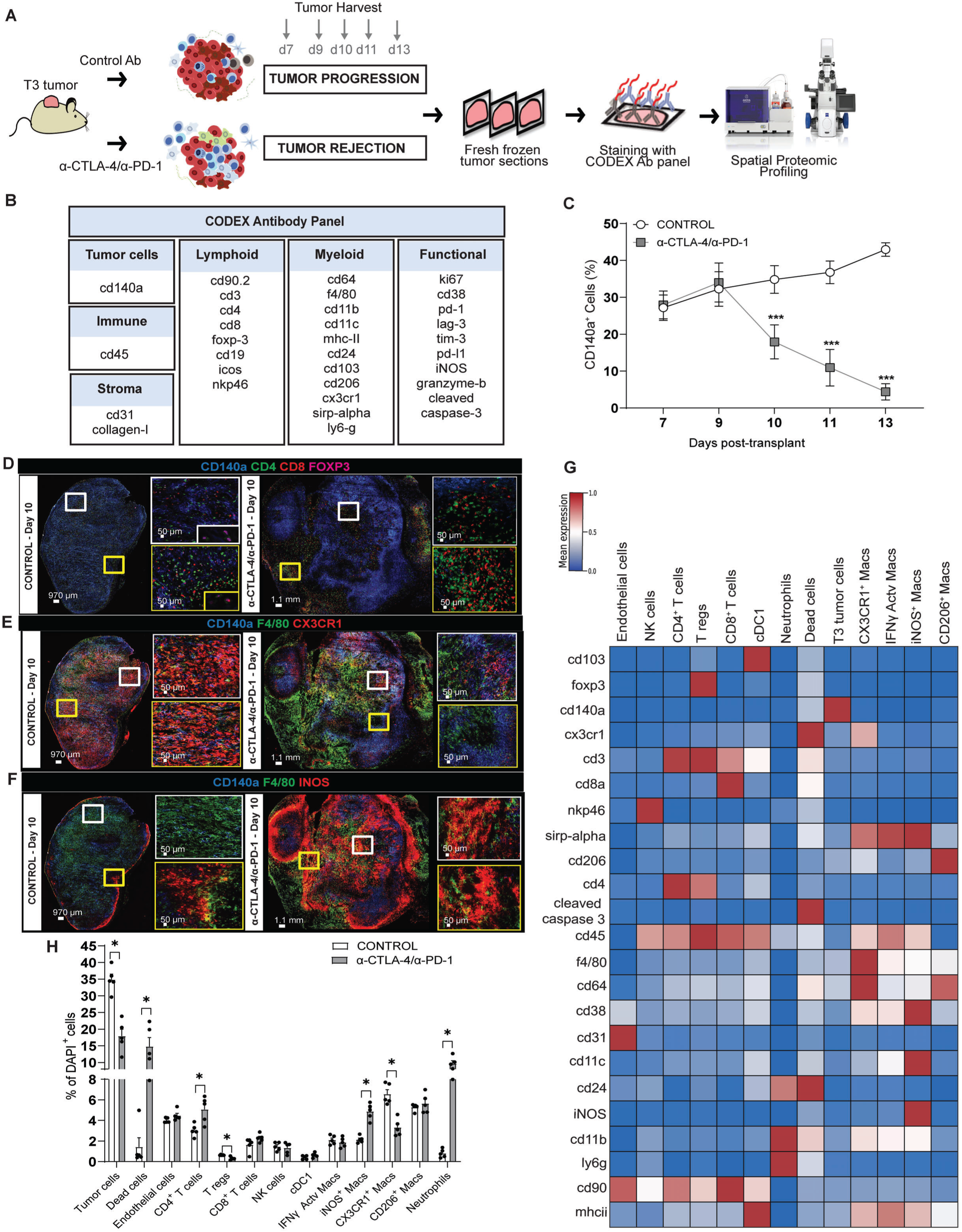
Spatial Proteomic Profiling to Monitor Successful Immune Checkpoint Therapy in T3 Tumors. **(A)** Schematic representation of the study experiment workflow. (**B**) CODEX antibody panel. **(C)** Longitudinal frequency of CD140a^+^ cells in T3 tumors from control mice [n=3(d7), n=4(d9), n=6(d10), n=5(d11), n=3(d13)] and α-CTLA-4/αPD-1 mice [n=3(d7), n=4(d9), n=5(d10), n=5(d11), n=4(d13)]. ***p=0.0002, ANOVA (with mixed-effects analysis) **(D)** D10 representative CODEX image with a four-color overlay showing CD140a^+^, CD4^+^, CD8^+^ and FOXP3^+^ cells. (**E)** D10 representative CODEX image with a three-color overlay for CD140a^+^, F4/80^+^ and CX3CR1^+^ cells. (**F)** D10 representative CODEX image with a three-color overlay for CD140a^+^, F4/80^+^ and iNOS^+^ cells. Scale bar as indicated. **(G)** Heatmap displaying normalized protein expression in each cell type. (**H)** Bar graph showing the cellular composition of T3 tumors harvested on d10. *p<0,05, Multiple unpaired t-test

DAPI^+^ cells were segmented, enabling us to obtain xy coordinates and single-cell protein marker intensity of 15,112,591 cells **(Figures S1C and S1D**). With this dataset, we visualized the longitudinal progression of CD140a^+^ tumor cells under control and ICT conditions (**Figure S1E)** and quantified their frequency over time to identify the earliest time point showing a decline in T3 tumor cell frequency following α-CTLA-4/α-PD-1 treatment **(Figure 1C)**. Samples from this initial time point were used to develop an analytical pipeline to assess spatial changes driving antitumor response. This analysis showed that the frequency of CD140a^+^ tumor cells remained stable on d7 and d9 post-transplant in both treatment groups but underwent a significant decrease on d10 where the percentage of tumor cells dropped from 34.8% in the control group to 17.9% in the α-CTLA-4/α-PD-1 group. By d13, the frequency of CD140a^+^ cells expanded in control mAb-treated mice to 42.9% but decreased to 4.3% in the α-CTLA-4/α-PD-1 group. This result indicated that ICT induces the rapid elimination of T3 tumors and highlights d10 as a critical time point marking the onset of therapeutic rejection under the treatment conditions employed.

In d10 tumors from control Ab-treated mice, CD4^+^, CD8^+^, and Foxp3^+^ T cells were found predominantly at the outer region of the tumor tissue. In contrast, α-CTLA-4/α-PD-1 induced pronounced intratumoral infiltration of CD4^+^ and CD8^+^ T cells **(Figure 1D)**. The myeloid compartment was also impacted by α-CTLA-4/α-PD-1. F4/80^+^CX3CR1^+^ macrophages were plentiful in tumors from control mice, but were reduced in numbers and localized to limited regions in the tumor after α-CTLA-4/α-PD-1 **(Figure 1E)**. Conversely, whereas iNOS^+^F4/80^+^ macrophages were only seen at the tumor border in control Ab-treated mice, they were broadly distributed in tumors from α-CTLA-4/α-PD-1 **(Figure 1F)**.

To confirm these morphological findings, we compared the results generated using immunocompetent wild-type mice to those obtained using syngeneic immunodeficient Rag2^-/-^mice, which lack T cells, B cells and NKT cells and thus fail to control tumor growth *in vivo*. As anticipated, α-CTLA-4/α-PD-1 did not induce T3 tumor rejection in *Rag2^-/-^* mice **(Figure S4A).** CODEX imaging of T3 tumors from Rag2^-/-^ mice revealed the absence of an ongoing anti-tumor response as evidenced by the absence of iNOS^+^ and CD38^+^ macrophages, suggesting a failure to elicit an *in vivo* IFNγ signature in the TME **(Figure S4B)**. Additionally, T3 tumor cells transplanted into *Rag2*^-/-^ mice grew more rapidly than those in wild-type mice, and the frequency of CD140a^+^ cells was not affected by α-CTLA-4/α-PD-1 in this immunodeficient background **(Figure S4C),** confirming the critical role of T cells in the tumor rejection process following α-CTLA-4/α-PD-1.

To profile our d10 dataset, we developed a hierarchical cell clustering approach that combines ANN classifiers, z-score normalization, and K-means over-clustering **(Figure S5A).** This method allowed the identification of 13 cell types, whose normalized protein expression values are displayed in **Figure 1G**. Dimensionality reduction via t-SNE confirmed the distinct identity of each immune cell populations **(Figure S5B and S5C),** which included: (i) 4 clusters of lymphoid cells: [CD3^+^CD4^+^ T cells; CD3^+^CD8^+^ T cells; NK cells (CD3^neg^NKP46^+^); and T regulatory cells (T-regs) (CD3^+^CD4^+^FOXP3^+^)]; (ii) conventional type 1 dendritic cells (cDC1), expressing MHCII^+^, CD11C^+^, CD103^+^, which in a separate experiment also co-stained for the XCR1 surface marker^27^ thus validating their identity **(Figure S5D);** (iii) neutrophils (expressing CD11B^+^LY6G^+^); and (iv) 4 populations of macrophage cells (identified as positive for CD64, CD11B, CD11C, F4/80 and SIRP-alpha), that consisted of CD206^+^ macrophages; CX3CR1^+^ macrophages; and two distinct IFNγ polarized macrophage population: one that expressed iNOS and CD38 [denoted iNOS^+^ macrophages (CD38^+^iNOS^+^)] and a second that expressed CD38 but lacked iNOS expression [denoted IFNγ activated macrophages) (CD38^+^iNOS^neg^)]. We previously showed that cytokines within the tumor microenvironment (TME) shape monocyte differentiation trajectories in T3 tumors^14^. These monocytes follow divergent differentiation trajectories, giving rise to either CX3CR1⁺ or iNOS⁺ macrophages. Intermediate phenotypic states along this continuum include CD38⁺iNOS⁻, and CD206⁺ macrophages. CD38 expression marks a macrophage phenotype driven by IFNγ, alone, while iNOS expression requires not only macrophage responses to IFN-γ but also responses driven by the NF-κB signaling pathway^14^ such as those induced by Tumor necrosis factor-alpha (TNF-α).

We also assessed changes in cellular composition at d10 following α-CTLA-4/α-PD-1 **(Figure 1H)**. Whereas the frequency of dead tumor cells in mice treated with control Ab was 1.4%, this frequency increased to 14.8% following α-CTLA-4/α-PD-1. Concomitantly, the proportion of CD4^+^ T cells at this time point increased from 3% in control to 5% with α-CTLA-4/α-PD-1. We also observed that the percentage of T-regs was markedly reduced by 50% (from 0.6% in control to 0.28% after α-CTLA-4/α-PD-1). The latter change was attributable to the known Treg-depleting activity of the α-CTLA-4 mAb (Clone 9D9) used in the α-CTLA-4/α-PD-1 mixture^28^. CD8^+^ T cells did not change significantly at d10 nor were differences detected in NK cells, cDC1, or endothelial cells. For other myeloid cells, we observed an expected increase in iNOS^+^ macrophages [from 2.1% (control) to 4.9% (α-CTLA-4/α-PD-1)]), a decrease in CX3CR1^+^ macrophages [from 6.6% (control) to 3.3% (α-CTLA-4/α-PD-1)], and an increase in neutrophils [0.8% (control) to 9.7% (α-CTLA-4/α-PD-1)]. These results recapitulate the remodeling of the lymphoid and myeloid populations that we had previously observed for α-CTLA-4/α-PD-1-treated, T3 tumor-bearing mice^14^ and demonstrate the suitability of our spatial proteomic profiling system for high-dimensional imaging of T3 tumors.

### *α*-CTLA-4/*α*-PD-1 Induces Co-Localization of CD4^+^ and CD8^+^ T Cells with Myeloid APCs in Day 10 Tumors

To define how α-CTLA-4/α-PD-1 alters spatial organization at the onset of tumor rejection, we characterized cell–cell co-localization at d10 using a permutation-based interaction analysis previously used by others ^29, 30^ **(Figure 2A)**. In tumors from mice treated with control mAb, only a minimal amount of co-localization was observed between CD4^+^ and CD8^+^ T cells. In contrast, following α-CTLA-4/α-PD-1, strong co-localization of CD4^+^ and CD8^+^ T cells (red box 1) was observed, and also between CD4^+^ T cells with cDC1 cells (red box 2) **(Figure 2B)**. We also observed that IFNγ-activated macrophages co-localized with CD4^+^ T cells post-α-CTLA-4/α-PD-1 (red box 3) **(Figure 2B)**. In the absence of α-CTLA-4/α-PD-1, Tregs co-localized with both CD4^+^ and CD8^+^ T cells (red box 4) and with cDC1 cells (red box 5) **(Figure 2B)**. α-CTLA-4/α-PD-1 reduced this co-localization, at least in part, because of αCTLA4-induced depletion of activated Tregs^28^.

**Figure 2.**
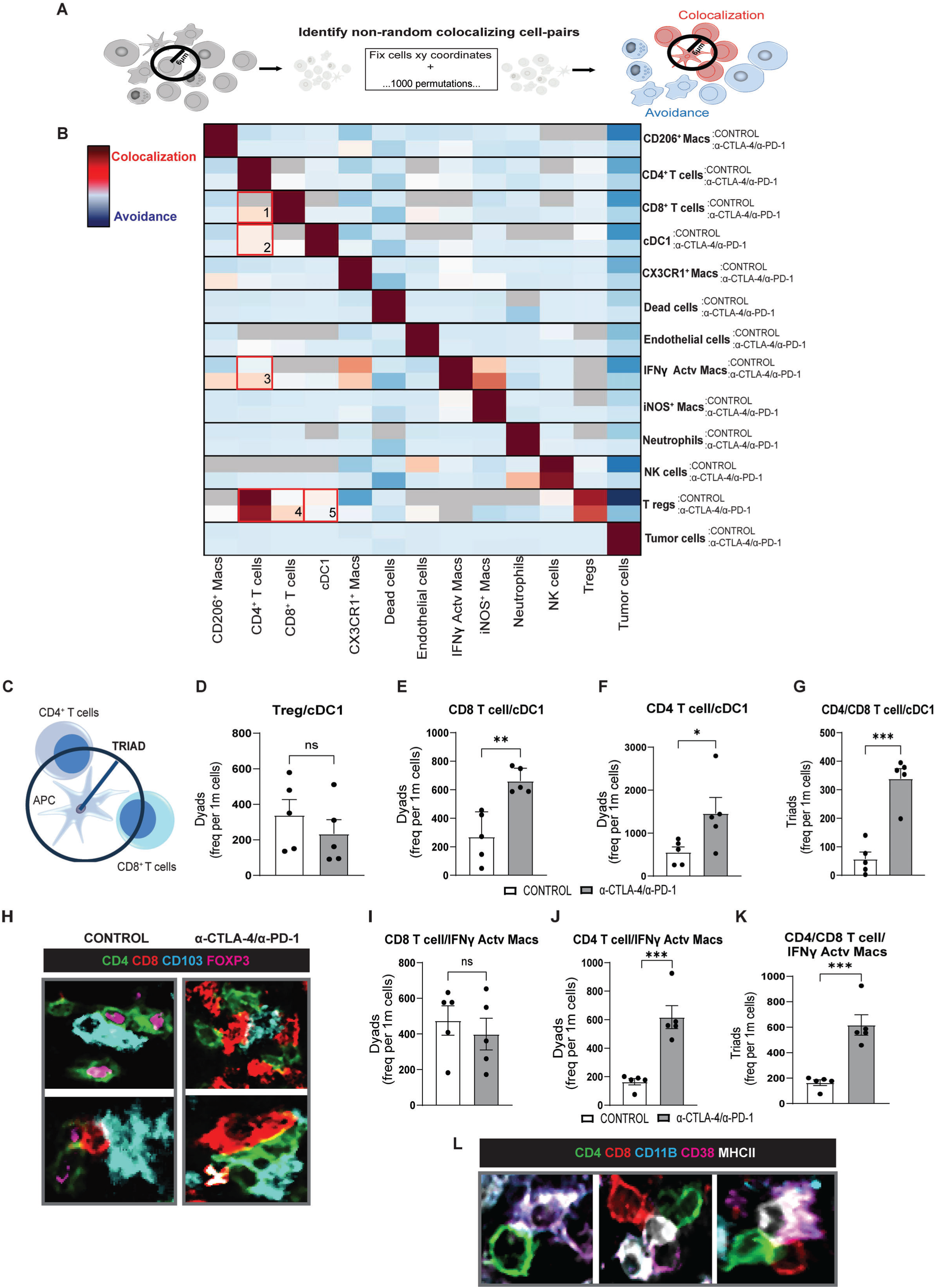
*α*-CTLA-4/*α*-PD-1 Induces Co-Localization of CD4^+^ and CD8^+^ T Cells with Myeloid APCs on Day 10. **(A)** Computational method used to identify non-random co-localizing cell pairs. A co-localization zone was defined as a 6 μm radius from the nucleus of a cell to the edge of a neighboring cell. Cells within this zone were considered co-localizing. To assess non-random co-localization, 1,000 label permutations were performed while keeping cell positions fixed. Significant co-localization was defined by a p-value < 0.01.**(B)** Heat map depicting significant cell-cell colocalization (red) or avoidance (blue) in control Ab (upper cell) or α-CTLA-4/α-PD-1 (lower cell). *p<0.05, adjusted p-value (Benjamini-Hochberg correction) after permutation test. (**C)** Schematic representation of the analysis used to quantify the frequency of triads between CD4^+^ and CD8^+^ T cells with myeloid APCs. (**D)** Frequency of dyads between Tregs and cDC1. **(E)** Frequency of dyads between CD8^+^ T cells and cDC1. **(F)** Frequency of dyads between CD4^+^ T cells and cDC1. (**G)** Frequency of triads between CD4^+^ T cells, CD8^+^ T cells and cDC1. **(H)** Four color overlay CODEX images displaying the cell interactions between CD4^+^ T cells, CD8^+^ T cells, Tregs (FOXP-3^+^) and cDC1 (CD103^+^) from control and α-CTLA-4/αPD-1 mice. **(I)** Frequency of dyads between CD8^+^ T cells and IFNγ-activated macrophages. **(J)** Frequency of dyads between CD4^+^ T cells and IFNγ-activated macrophages. **(K)** Frequency of triads between CD4^+^ T cells, CD8^+^ T cells and IFNγ-activated macrophages. **(L)** Five color overlay CODEX images displaying the cell interactions between CD4^+^ T cells, CD8^+^ T cells and IFNγ-activated macrophages (CD11B^+^CD38^+^MHC-II^+^). ns= non-significant.*p=0.0496,**p=0.0021,***p=0.0002, Unpaired t-test. From B to K, n=5 for control and n=5 for α-CTLA-4/αPD-1.

This analysis identified CD4^+^ T cells, CD8^+^ T cells, cDC1 and IFNγ-activated macrophages as immune cells whose co-localization in the tumor are positively impacted by α-CTLA-4/α-PD-1 treatment on d10. Given the recently established role of three-cell clusters (triads), in which CD4⁺ T cells co-engage with CD8⁺ T cells and APCs to present both CD8⁺ and CD4⁺ tumor antigens and effectively license CD8⁺ T cell cytotoxicity^31^, we further investigated how the cellular partnership of cDC1 and IFNγ-activated macrophages changed following α-CTLA-4/α-PD-1 **(Figure 2C).** We initially assessed the frequency by which cDC1 formed either two-cell clusters (dyads) with Tregs, CD4⁺ or CD8⁺ T cells, or triads consisting of an individual cDC1 interacting with both a CD4⁺ and a CD8⁺ T cell. Treg-cDC1 dyads were found in tumors derived from either control mAb or α-CTLA-4/α-PD-1 mice, and did not show significant differences between the mAb conditions used **(Figure 2D**, p=0.4). In contrast, following α-CTLA-4/α-PD-1, cDC1 were more frequently observed in dyads with CD8^+^ T cells (**Figure 2E**, p=0.002) or CD4^+^ T cells **(Figure 2F**, p=0.004) compared to control. α-CTLA-4/α-PD-1 also induced a highly significant increase in the frequency of cDC1-CD4-CD8 triads **(Figure 2G**, p=0.0002). Representative images of these cellular interactions are presented in **Figure 2H**.

We then performed the same analysis focused on IFNγ-activated macrophages. Here, the frequency of dyads between IFNγ-activated macrophages and CD8^+^ T cells did not change with or without α-CTLA-4/α-PD-1 (**Figure 2I**, p=0.88). However, we observed a significant increase in the frequency of dyads containing CD4^+^ T cells following α-CTLA-4/α-PD-1 (**Figure 2J**, p=0.03). Similarly, triads between IFNγ-activated macrophages and both CD4^+^ and CD8^+^ T cells increased after α-CTLA-4/α-PD-1 (**Figure 2K**, p=0.0006). Images of the cellular interactions are presented in **Figure 2L**. The detection of these immune triads suggests that α-CTLA-4/α-PD-1 promotes coordinated multicellular interactions between T cells with two professional APCs in the tumor: cDC1s and IFNγ-activated macrophages. Even though cDC1s have a known superior ability to *prime* naïve T cells in the lymph node, our results also support a potential role for IFNγ-activated macrophages in the tumor for supporting the effector phase of the anti-tumor response through interactions with CD4^+^ and CD8^+^ T cells.

### Neighborhood Mapping Reveals Coordinated Redistribution of Immune Cells Leading to the Formation of Seven Cellular Neighborhoods (CNs) in T3 Tumors

We reasoned that pairwise cell-cell interactions represented only the first step in defining the spatial considerations by which anti-tumor immune responses are organized. To gain a more comprehensive view of higher-order tissue organization, we proceeded to identify multicellular networks within the tumor tissue, often referred to as “Cellular Neighborhoods” (CNs)^18^. In particular, we were interested in assessing whether α-CTLA-4/α-PD-1 was associated with a neighborhood formed by increased interactions between T cells, cDC1, and IFNγ-activated macrophages. To achieve this, we employed a ‘Sliding Window Mapping’^18^ strategy to identify prominent CNs ^18^ **(Figure 3A)**.

**Figure 3.**
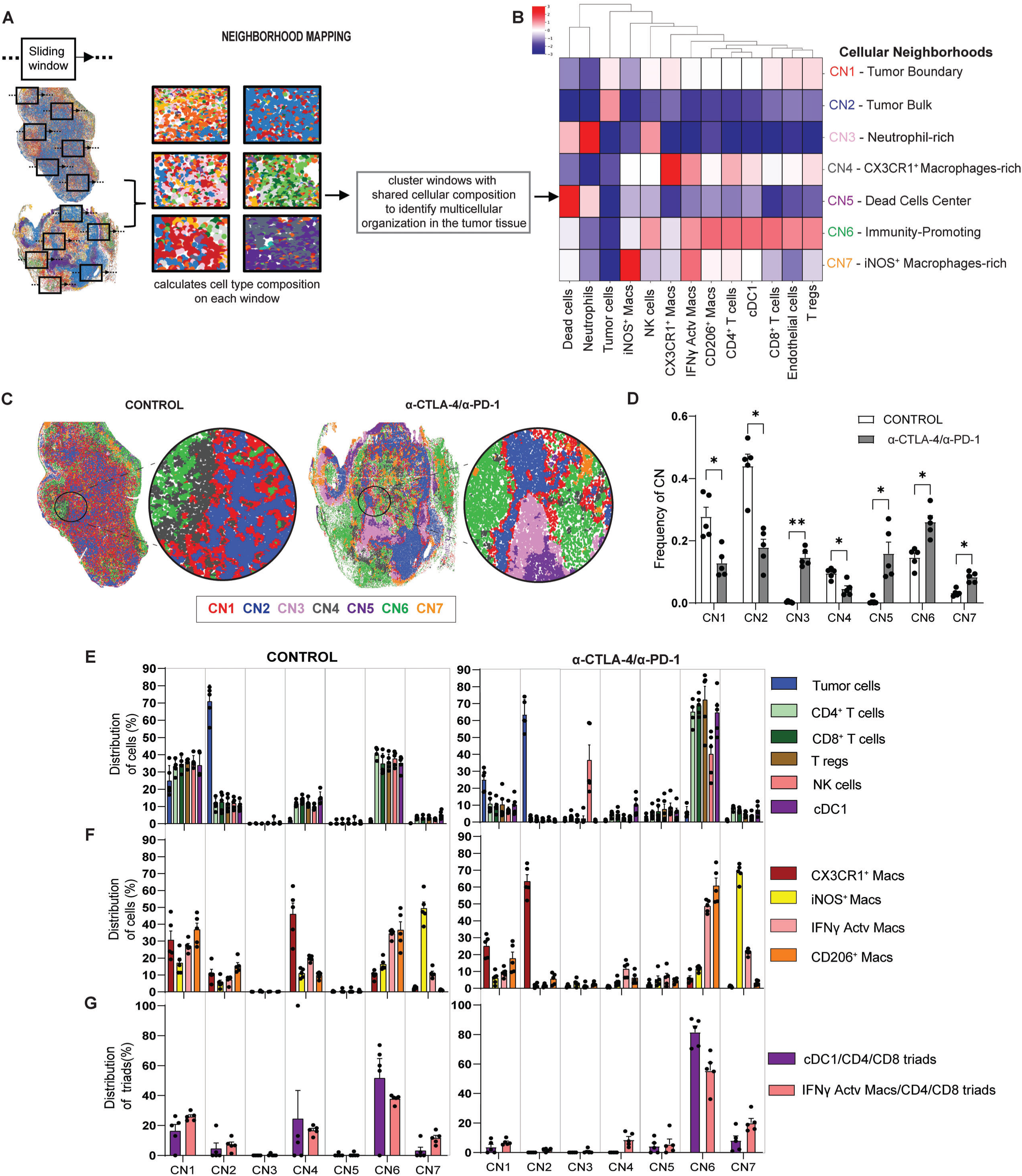
Neighborhood Mapping Reveals Coordinated Redistribution of Immune Cells Leading to the Formation of Seven Cellular Neighborhoods (CNs) in T3 Tumors. **(A)** Schematic representation of the Neighborhood Window Mapping method in T3 tumors harvested on d10. This method works iteratively by forming a capture window around each cell, profiling its nearest neighbors, and using the cellular composition of each window to identify the different CNs. (**B**) Heat map showing the identification of seven distinct CNs in T3 tumors and their respective frequencies (enrichment score) within each CN (pooled data from both groups). n=5 control and n=5 α-CTLA-4/αPD-1. (**C)** Representative dot plots depicting the seven CNs and their corresponding colors. (**D)** Frequency of CNs in d10 T3 tumors from control Ab versus α-CTLA-4/αPD-1 mice. ***p < 0.001, T-test. **(E)** Percentage of Tumor cells, CD4^+^ and CD8^+^ T cells, Tregs, NK cells, and cDC1 distributed across the CNs in d10 T3 tumors. **(F)** Percentage of CX3CR1^+^ macrophages, iNOS^+^ macrophages, IFNγ activated macrophages, and CD206^+^ macrophages distributed across CNs in d10 T3 tumors. **(G)** Percentage of cDC1 and IFNγ activated macrophage triad cells distributed across the CNs in d10 T3 tumors. From D to G, n=5 for control and n=5 for α-CTLA-4/αPD-1.

With this approach, we identified seven distinct CNs in T3 tumors from both control and α-CTLA-4/α-PD-1 mice **(Figure 3B)**. The detailed cellular composition of each CN is shown in **Figure S6A**. CN1 comprises the “Tumor Boundary” and consists of 47.7% CD140a^+^ tumor cells and a mixture of immune cells that includes macrophages and T cells (red color in **Figure 3C**). CN2 is composed of 90% CD140a^+^ tumor cells and thus was named “Tumor Bulk” (blue color in **Figure 3C**). CN3 consisted mostly of neutrophils (70%) and dead cells (16%) and was named “Neutrophil-rich” (light magenta color in **Figure 3C**). CN4 represents the neighborhood with the highest concentration of CX3CR1^+^ macrophages (47% of this CN composition) and was named “CX3CR1^+^ macrophage-rich” (dark gray in **Figure 3C**). CN5 denotes the “Dead Cell Center” (dark purple in **Figure 3C**) since it contained 72% dead cells and resides in the inner region of CN3. CN6 displayed the highest enrichment of CD4^+^ and CD8^+^ T cells compared to the other CNs (i.e., CD4^+^ and CD8^+^ T cells comprise 17% and 8.4% of CN6, respectively). Importantly, CN6 also contained a mixture of NK cells (2.9%), endothelial cells (17%), and different populations of myeloid cells, such as cDC1 (2%), CD206^+^ macrophages (18%), and IFNγ-activated macrophages (4%). CN6 was labeled as “Immunity-Promoting” (hereafter IP-CN6) because it was comprised of key, spatially organized antitumor lymphoid populations that colocalized with different types of myeloid APCs. Lastly, CN7 was labeled “iNOS^+^ macrophage-rich” as it contains 55% of iNOS^+^CD38^neg^ and 6.5% of iNOS^neg^CD38^+^ macrophages (yellow color in **Figure 3C**).

As shown in **Figure 3D**, α-CTLA-4/α-PD-1 induced significant changes in the frequencies of all seven CNs. Specifically, the frequencies of CN1 (tumor boundary), CN2 (tumor bulk) and CN4 (CX3CR1^+^ macrophage-rich) were reduced in tumors from α-CTLA-4/α-PD-1 mice. In contrast, α-CTLA-4/α-PD-1 induced an increase in IP-CN6. We also noted increases in neutrophil-rich (CN3), dead cell center (CN5), and iNOS^+^ macrophage-rich (CN7) CNs. Collectively, these changes suggest that α-CTLA-4/α-PD-1 remodels immune cell networks such that they might potentially favor tumor elimination.

While this analysis identified changes in tissue organization, it did not address how α-CTLA-4/α-PD-1 affects the distribution of individual cell types within the CNs. To investigate this, we quantified the percentage of each cell type across CNs in tumors from control and α-CTLA-4/α-PD-1-treated mice at d10 **(Figure 3E)**. In control mAb-treated mice, 75% of CD140a^+^ tumor cells were localized within the tumor bulk (CN2), with the remaining 35% found at the tumor boundary (CN1). This distribution pattern remained unaltered upon α-CTLA-4/α-PD-1, since the majority of tumor cells continued to occupy CN2 and CN1. However, α-CTLA-4/α-PD-1 induced a marked shift in the distribution of immune cells. In tumors from control mAb-treated mice, approximately 35% of T cells were present at the tumor boundary (CN1) and similarly at IP-CN6. The remaining CD4^+^ and CD8^+^ T cells were distributed between the tumor bulk CN2 (12%) and at the CX3CR1^+^ macrophage-rich region CN4 (∼13%) **(Figure 3E)**. In contrast, tumors from α-CTLA-4/α-PD-1 mice exhibited a striking accumulation of CD4^+^ T cells (65%) and CD8^+^ T cells (68%) within IP-CN6. Notably, while the overall frequency of CD8^+^ T cells in the tumor remained unchanged at d10, their redistribution into IP-CN6 increased almost 2-fold to 68%. Additionally, the Tregs that were not depleted by α-CTLA-4/α-PD-1 remained localized in IP-CN6, although they represented only 1% of the IP-CN6 cellular population **(Figure 3E)**. NK cells clustered in IP-CN6 and CN3.

We used a similar approach to define the distribution of the different myeloid cell populations. In tumors from control mAb-treated mice, CD38^+^iNOS^neg^ and CD206^+^ macrophages accumulated at the tumor boundary (CN1) (27%) and in IP-CN6 (34%) but showed a sharp increased accumulation into IP-CN6 after α-CTLA-4/α-PD-1 (48%) **(Figure 3F)**. In contrast, CX3CR1^+^ macrophages displayed a distinct distribution pattern. In tumors from control mice, 45% of CX3CR1^+^ macrophages were located in CN4 and at the CN1 tumor boundary (30%) **(Figure 3F)**. After α-CTLA-4/α-PD-1, most of the CX3CR1^+^ macrophages became undetectable in CN4 (0.9%), and the few that remained were mostly found in the tumor bulk CN2 (63%). iNOS^+^ macrophages increased in CN7 from 49% to 68% after α-CTLA-4/α-PD-1 **(Figure 3F)**.

In addition, we noted an alteration in the distribution of neutrophils and dead cells. In tumors from control mice, both cell types were found dispersed across all seven CN but mostly accumulating in IP-CN6, with 25% of neutrophils and 37% of dead cells located in this CN **(Figure S6B).** Following α-CTLA-4/α-PD-1, the concentration of these cell types decreased in IP-CN6 and increased in their respective CNs. Specifically, 77% of the neutrophils localized in CN3, while 55% of dead cells were found in the dead cell center CN5 **(Figure S6B)**. This indicates that while there is a substantial increase in the frequency of neutrophils and dead cells after α-CTLA-4/α-PD-1, their location is restricted to specific areas of the tumor tissue and does not impact the composition of IP-CN6.

The accumulation of T cells and myeloid cells within IP-CN6 after α-CTLA-4/α-PD-1 suggests that this CN serves as a preferential site for immune interactions. To confirm this, we assessed the proportion of cDC1s and IFNγ-activated macrophages forming triads with CD4⁺ and CD8⁺ T cells (previously identified in **Figures 2G** and **2K**, respectively) across the seven CNs **(Figure 3G).** In tumors from control mice, 56% of cDC1 and 39% of IFNγ-activated macrophage triad-forming cells were in IP-CN6, and the remainder was shared between the tumor boundary CN1, tumor bulk CN2, and CX3CR1^+^macrophage-rich CN4. However, following α-CTLA-4/α-PD-1, we observed a very pronounced accumulation of triad-associated cDC1s (90%) and IFNγ-activated macrophages (56%) specifically within IP-CN6. This spatial enrichment of immune triads supports the role of IP-CN6 as a site for productive interactions between T cells and myeloid APCs.

Overall, these findings show a profound reorganization of the T3 TME, mediated by a coordinated redistribution of multiple immune cell types. α-CTLA-4/α-PD-1 thus induces a focused response from CD4^+^ and CD8^+^ T cells that, together with cDC1, CD206^+^ macrophages, and IFNγ-activated macrophages, drive the formation of IP-CN6.

### TNF*α* and IFN*γ* Signaling are Putative Communication Pathways Between T cells and Myeloid APCs within IP-CN6

While CODEX enables spatial proteomic profiling, it does not allow unbiased discovery of the molecular pathways underlying the formation and function of IP-CN6. To address this limitation, we generated paired scRNA-seq data from immune and tumor cells isolated from T3 tumors harvested on d10 (**Figure S7A and S7B**), and integrated this data with our d10 CODEX dataset using ARCADIA (ARchetype-based Clustering and Alignment with Dual Integrative Autoencoders)^32^ **(Figure 4A)**. ARCADIA is a generative framework for integrating weakly linked single-cell data modalities without requiring feature-level correspondence or paired cell barcodes, thus enabling spatially informed analysis of transcriptional programs.^32^. Specifically, it establishes cross-modal correspondence by first identifying modality-specific archetypes that define distinct cellular states and then aligning them across modalities based on cell-type distribution^32^ (**Figure S7C and S7D**). This alignment yields an entangled embedding that preserves cell type composition as well as spatial neighborhood structure, effectively transferring spatial context from CODEX to cells profiled with scRNA-seq and enabling CN-specific differential gene expression (DEG) analysis.

**Figure 4.**
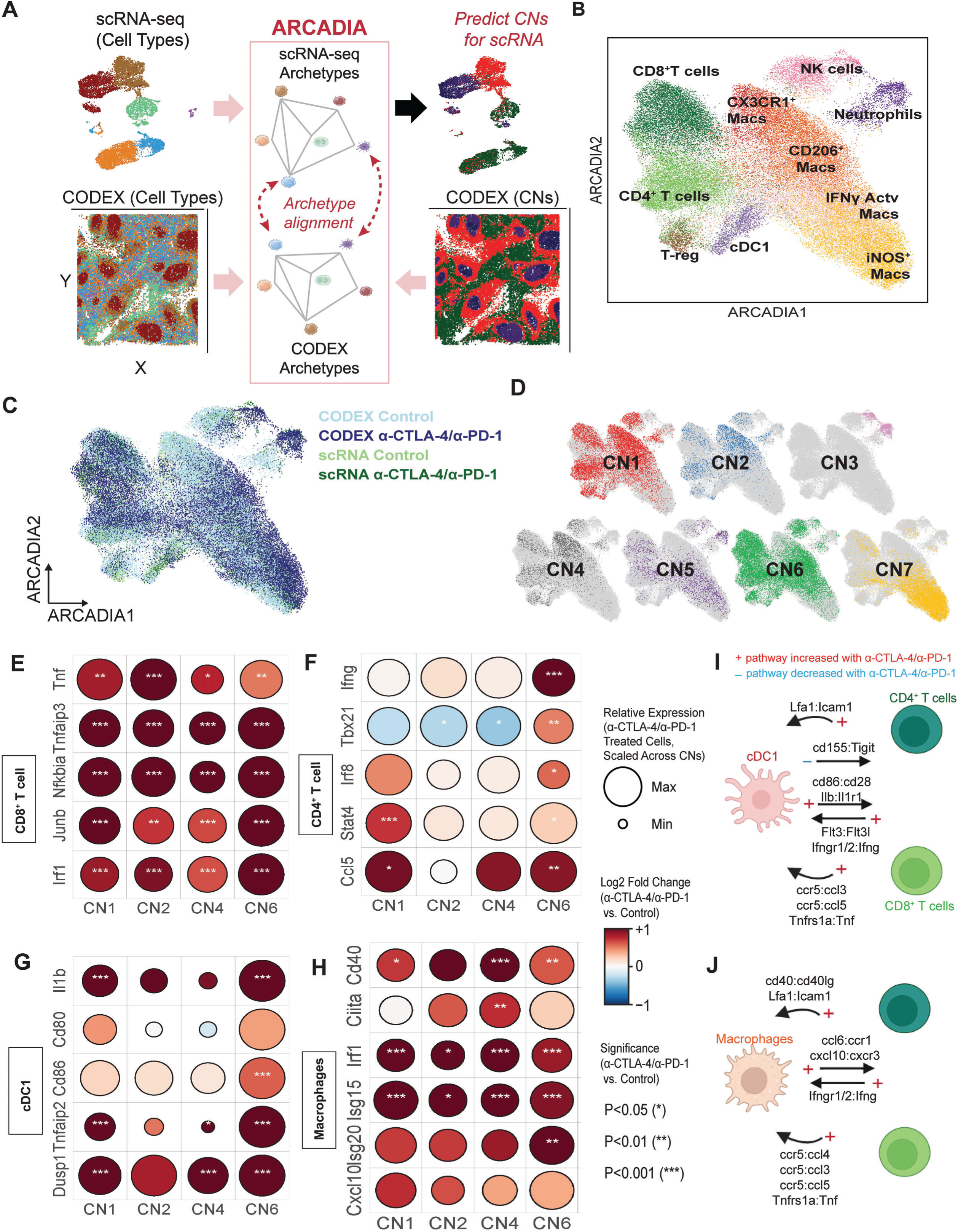
TNF*α* and IFN*γ* Signaling are Putative Communication Pathways Between T cells and Myeloid APCs within IP-CN6. **(A)** Overview of ARCADIA framework for integrating scRNA-seq and CODEX data, including input and output information. **(B,C)** ARCADIA integrated latent embedding colored by immune cell type (B) and modality and treatment condition (C). **(D)** UMAP of cells colored by known (ground truth) CNs in CODEX profiled cells. **(E-H)** Differentially expressed genes (DEGs) between α-CTLA-4/α-PD-1 and control conditions for CD8^+^ T **(E)**, CD4^+^ T **(F)**, cDC1 **(G)**, and macrophage **(H)** cells across CNs. Dot size corresponds to mean expression of each marker for α-CTLA-4/α-PD-1-treated cells within a CN, and sizes are scaled for each marker across CN groups. Log_2_ fold change and significance for each dot represent differential expression comparisons for each marker between α-CTLA-4/α-PD-1-treated and control cells within a CN **(I)** Schematic figure summarizing cell-cell communication pathways between CD4^+^ T cells, CD8^+^ T cells and cDC1 from ARCADIA and ContactTracing analyses. **(J)** Schematic figure summarizing cell-cell communication pathways between CD4^+^ T cells, CD8^+^ T cells and macrophages from ARCADIA and ContactTracing analyses. Detailed circus plots are depicted in **Figure S9** and statistical analysis in **Table S3**.

When applied to our d10 scRNA and spatial protein data, ARCADIA demonstrated consistent archetype representations across modalities **(Figure S7E)** and accurate representation of cell types in the shared embedding **(Figure 4B and 4C)**. Importantly, ARCADIA-enabled inference of CN labels for scRNA-seq cells recapitulated CN composition observed in CODEX data (**Figure S8A**), including CN1, CN2, CN4, and IP-CN6, as evidenced by clustering according to CN identity in the integrated embedding **(Figure 4D)**. Since ARCADIA relies on shared cell-type representation across modalities for archetype alignment, downstream analyses were restricted to CNs with sufficient cellular representation, excluding neutrophils and dead cells.

We next performed Gene Set Enrichment Analysis (GSEA) on CD8⁺ T cells assigned to IP-CN6 to identify molecular programs enriched within this neighborhood. Following α-CTLA-4/α-PD-1. treatment, CD8⁺ T cells exhibited significant enhancement of TNFα expression as well as its downstream NF-κB signaling pathways (**Figure S8B**), accompanied by increased expression of key target genes, including *A20* (*Tnfaip3)*, *Junb*, and *Irf1*. This signature was also observed in CN1, CN2 and CN4, indicating broadly enhanced effector activation across multiple neighborhoods **(Figure 4E)**. In contrast, GSEA of CD4⁺ T cells revealed significant enrichment of the IFNγ response pathway after α-CTLA-4/α-PD-1 **(Figure S8C)**, consistent with a Th1 phenotype. This was particularly pronounced within IP-CN6 and was characterized by increased expression of *IFNγ*, *Tbx21* (T-bet), *Stat4*, *Irf8*, and *Ccl5* **(Figure 4F)**. These findings suggest that CD4⁺ T cells within IP-CN6 adopt a helper phenotype that supports CD8⁺ T cell effector function. Consistent with this inflammatory milieu, cDC1s exhibited enrichment of TNFα signaling components and increased expression of inflammatory and antigen-presentation genes, including *Il-1b (Interleukin-1 beta)*, *B94* (*Tnfaip2)* and *Cd86*, indicative of enhanced maturation and co-stimulatory capacity (**Figure 4G)**. Similarly, macrophages within IP-CN6 displayed enrichment of type I and type II IFN response pathways as reflected by increased expression of *Cd40*, *Irf1*, Isg15, and Isg20 **(Figure 4H)**.

To further validate these intercellular communication pathways, we performed ContactTracing-based ligand–receptor analysis^33^, which revealed extensive remodeling of signaling networks following α-CTLA-4/α-PD-1 treatment **(Fig. S9A and S9b; Table S3, summarized in Fig. 4I–J)**. This analysis predicted TNFα signaling from CD8⁺ T cells to both cDC1s **(Fig. 4I)** and macrophages **(Fig 4J)**, which were shown to express TNFR1 (Tnfrsf1a). Similarly, increased levels of IFNγ were predicted to affect both cDC1s and macrophages via signaling through functional IFNγ receptors (Ifngr1/Ifngr2). Changes in chemotactic signatures were also observed, as shown by an increase in CCL3 and CCL4 expression by CD8⁺ T cells, together with CCL5 derived from both CD4⁺ and CD8⁺ T cells, which can recruit myeloid cells via CCR5 **(Fig. 4I-J)**. Conversely, macrophage-derived CXCL10 and CCL6 were found to communicate with CD4⁺ and CD8⁺ T cells, suggesting that multiple chemotactic pathways promote interactions between lymphoid and myeloid populations **(Fig. 4J).** Consistent with enhanced T cell activation, the CD86:CD28 co-stimulatory pathway from cDC1s to T cells was significantly increased **(Fig. 4I).** Additional pathways associated with APC activation and expansion were also enriched, including Flt3Lg (Fms-related tyrosine kinase 3 ligand) derived from T cells, which is known to promote cDC1 differentiation and survival^34^ **(Fig. 4I)**. Notably, the CD40L: CD40 costimulatory axis between CD4⁺ T cells and macrophages was enhanced **(Fig. 4J)**, which is a well-established pathway for CD4^+^ T cells to trigger robust macrophage activation^35^. Communication between cell-contact molecules, such as Icam1 (intercellular adhesion molecule-1) and its receptor Lfa1 (leukocyte function-associated antigen-1), were also positively impacted, supporting improved cell-to-cell interactions. Lastly, communication between the inhibitory checkpoint molecule TIGIT (T cell immunoreceptor with Ig and ITIM domains) and its receptor CD155 was reduced **(Fig. 4I)**. Although this analysis cannot distinguish signaling occurring specifically within triad versus dyad interactions, the spatially informed transcriptional programs inferred by ARCADIA highlight plausible pathways that are likely to contribute to the assembly and function of IP-CN6 during α-CTLA-4/α-PD-1 therapy.

### Longitudinal Analysis Reveals Cellular Dynamics Driving the Expansion of the Immunity-Promoting CN6 During Tumor Rejection

Thus far, our analysis revealed the spatial and molecular changes that occur at d10, a time point at which the α-CTLA-4/α-PD-1-induced rejection process was just beginning. To understand the longitudinal evolution of the seven CNs, in particular IP-CN6, we applied our hierarchical analysis pipeline **(Figure S5A)** to a dataset encompassing tumors harvested on d7, d9, d10, d11, and d13 post-tumor transplant.

This longitudinal analysis revealed the dynamics of CD4^+^ and CD8^+^ T cells **(Figure 5A)**. In tumors from control mAb-treated mice CD4^+^ T cells accumulated until d9, reaching 3.6% of total cell composition, but then steadily declined thereafter to 1.1% on d13 **(Figure 5B)**. In contrast, tumors from α-CTLA-4/α-PD-1 mice displayed the opposite CD4^+^ T cell trajectory. Specifically, α-CTLA-4/α-PD-1 induced an increase in CD4^+^ T cells over control that first became detectable at d10 and continued to rise through d13 to 6.2%. This dynamic coincided with the elimination of CD140a^+^ tumor cells **(Figure 1C)**. Intriguingly, the increase in CD4^+^ T cells following α-CTLA-4/α-PD-1 preceded a significant rise in CD8^+^ T cells, which was detected on d11. CD8^+^ T cells remained below 2% until d11 but then surged in tumors from α-CTLA-4/α-PD-1-treated mice, peaking at 7.5% on d13, eventually surpassing the frequency of CD4^+^ T cells **(Figure 5C)**. NK cell frequency did not change significantly at any time point **(Figure S10A),** while, as expected, Treg frequency remained significantly higher in tumors from control mice compared to mice receiving α-CTLA-4/α-PD-1 **(Figure S10B)**.

**Figure 5.**
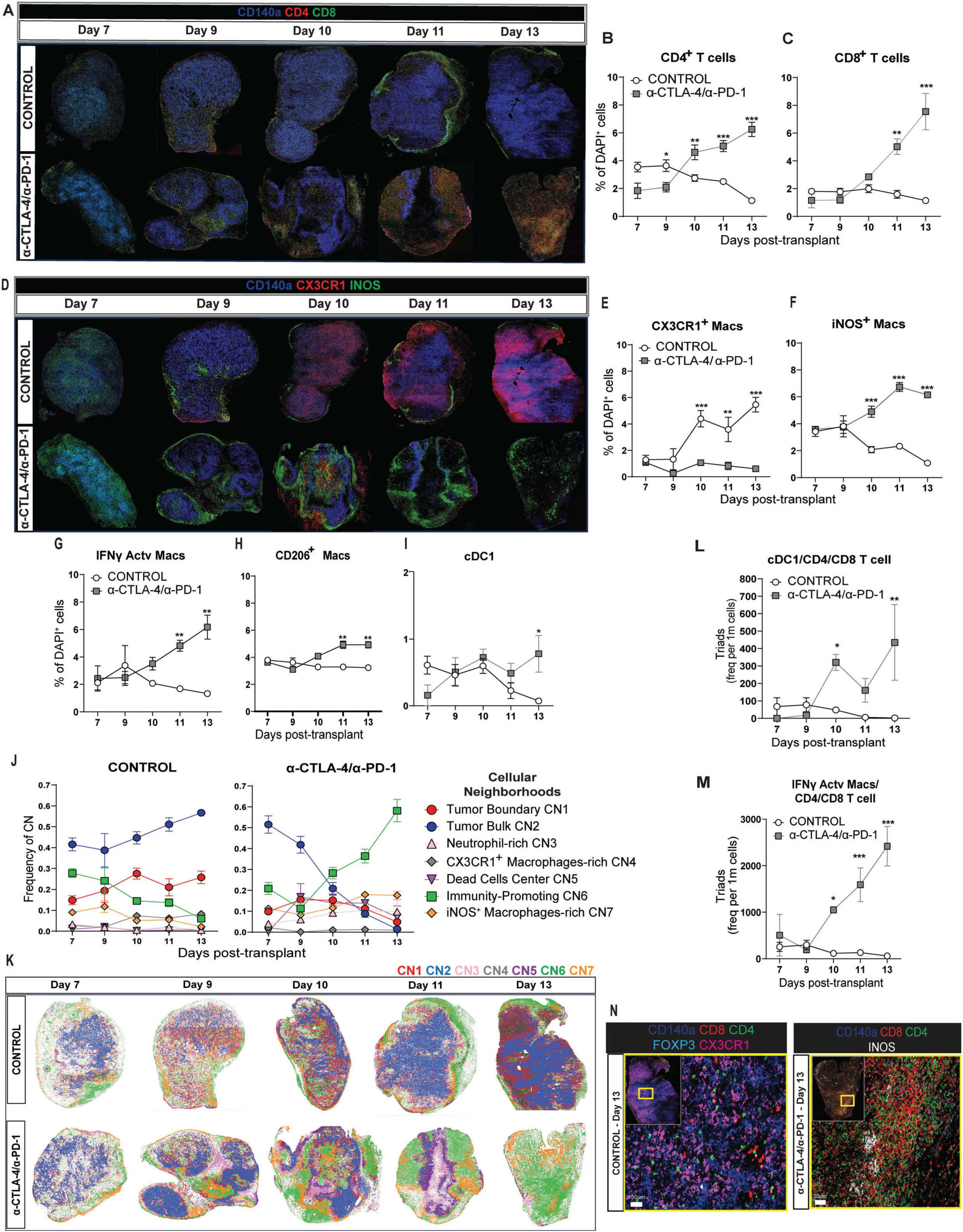
Longitudinal Analysis Reveals Cellular Dynamics Driving the Expansion of the Immunity-Promoting CN6 During Tumor Rejection. **(A)** Longitudinal three-color CODEX images of CD140a^+^, CD4^+,^ and CD8^+^ cells of T3 tumors harvested in multiple time points from control Ab and α-CTLA-4/α-PD-1 mice. **(B)** Longitudinal frequency of CD4^+^ T cells. **p=0.0079, ****p<0.0001, ANOVA (with mixed-effects analysis). **(C)** Longitudinal frequency of CD8^+^ T cells.***p<0.0001, ANOVA (with mixed-effects analysis). **(D)** Longitudinal three-color CODEX images of CD140a^+^, iNOS^+,^ and CX3CR1^+^ cells of T3 tumors harvested at multiple time points from control Ab and α-CTLA-4/α-PD-1 mice. **(E)** Longitudinal frequency of CX3CR1^+^ macrophages. **p=0.0014, ***p<0.0001, ANOVA (with mixed-effects analysis). **(F)** Longitudinal frequency of iNOS^+^ macrophages. ***p<0.0001, ANOVA (with mixed-effects analysis). **(G)**Longitudinal frequency of IFNγ-activated macs. **p=0.0012, ANOVA (with mixed-effects analysis). **(H)** Longitudinal frequency of CD206^+^ macrophages. **p=0.0019, ANOVA (with mixed-effects analysis). **(I)** Longitudinal frequency of cDC1. *p<0.05, ANOVA (with mixed-effects analysis). **(J)** Longitudinal frequency analysis of the seven CNs in T3 tumors from control Ab and α-CTLA-4/αPD-1 mice. **(K)** Representative dot plots depicting the seven CNs and their corresponding colors in the longitudinal dataset. **(L)** Longitudinal frequency of triads between CD4^+^ T cells, CD8^+^ T cells and cDC1. **(M)** Longitudinal frequency of triads between CD4^+^ T cells, CD8^+^ T cells and IFNγ-activated macrophages. **(N)** Representative high magnification CODEX image of a T3 tumor harvested on d13. Scale bar is 50μm. From B to M, mean in Control represents [n=3(d7), n=4(d9), n=6(d10), n=5(d11), n=3(d13)] and in α-CTLA-4/αPD-1 [n=3(d7), n=4(d9), n=5(d10), n=5(d11), n=4(d13)].

This longitudinal analysis also revealed changes in the myeloid compartment **(Figure 5D)**. CX3CR1^+^ macrophages were first detected on d10 in mice treated with control Ab and became the major immune cell population by d13, reaching 5.4% of total cells **(Figure 5E**). In contrast, following α-CTLA-4/α-PD-1, CX3CR1^+^ macrophages became nearly undetectable while iNOS^+^ macrophages and IFNγ activated macrophages (CD38^+^iNOS^neg^) increased to 6.1% each **(Figure 5F and 5G**, respectively). By d13, CD206^+^ macrophages represented 3.2% in tumors from control mice and 4.9% in α-CTLA-4/α-PD-1-treated mice **(Figure 5H)**. cDC1 frequency remained unchanged until d13, when it dropped in tumors from control mice while increasing in tumors from α-CTLA-4/α-PD-1 mice (p=0.04) **(Figure 5I)**. Lastly, we noted that in rejecting tumors, both dead cells and neutrophils had a transitory dynamic **(Figure S10C)**. Neutrophils were only significantly increased on d10 (p=0.0008), a day after dead cells increased, and lost significance thereafter by d13 (p=0.4) **(Figure S10D)**. Dead cells significantly increased on d9 (p=0.0008), reaching 11.6%, but then fell to only 3.1% by d13, losing significance (p=0.2) **(Figure S10E)**.

To analyze how these cellular dynamics impacted the T3 tumor tissue architecture temporally, we pooled the data from all time points and applied the sliding window mapping method to identify the CNs. The same seven CNs identified on d10 were consistently observed across different stages of tumor growth or regression, although at differing relative magnitudes, indicating that these structures are quasi-stable but are dynamically influenced during the organization and destruction of T3 tumors **(Figure S10F)**. In tumors growing progressively in control mAb-treated mice, the tumor boundary (CN1) and tumor bulk (CN2) became the largest CNs by d13 **(Figure 5J)**. IP-CN6 initially emerged as the second-largest neighborhood, but its frequency then declined over time. Other CNs showed only minimal changes and also decreased over time. In contrast, tumors from α-CTLA-4/α-PD-1 mice showed a marked longitudinal reduction in tumor boundary (CN1) and tumor bulk (CN2) due to the destruction of tumor cells. In contrast, IP-CN6 steadily expanded from d10 on, eventually becoming the largest CN in rejecting tumors by d13 **(Figure 5J)**. The iNOS^+^ macrophage-rich CN7 developed as the second largest CN post α-CTLA-4/α-PD-1, followed by the neutrophil-rich CN3 region. The longitudinal evolution of the 7 CNs is presented in **Figure 5K**.

Given the role of IP-CN6 as a site for immune interactions, we next analyzed the longitudinal dynamics of immune triads involving T cells with cDC1s or IFNγ–activated macrophages **(Figures 5L and 5M**, respectively). In tumors from control Ab-treated mice, the frequency of triads nucleated by both types of APCs declined over time and became undetectable by d13. However, tumors from α-CTLA-4/α-PD-1-treated mice showed a progressive increase in three-cell cluster interactions, starting at d10 and peaking at d13. This increase was more pronounced for triads involving IFNγ–activated macrophages, likely reflecting their higher abundance in the T3 TME compared to cDC1s. Nevertheless, the timing and evolution of these clusters between APCs and T cells coincided with the amplification of the anti-tumor immune response.

These results highlight the emergence of two distinct TMEs by d13: one associated with progressive tumor growth and the other associated with effective immune-mediated tumor rejection **(Figure 5N).** In mice treated with control mAb, the cellular components of IP-CN6 declined over time, including the aforementioned triads, while CX3CR1⁺ macrophages and tumor cells expanded, driving the growth of CN4 and CN2, respectively **(Figure 5N)**. In contrast, α-CTLA-4/α-PD-1 led to a rapid expansion of CD4⁺ and CD8⁺ T cells, while sustaining the number of triads with cDC1 and IFNγ–activated macrophages, resulting in the continued expansion of the IP-CN6.

### Granzyme B*⁺* and Ki67*⁺* T Cells Form Distinct Sub-Neighborhoods (IP-SNs) Within Immunity-Promoting CN6

Our findings thus far suggested that IP-CN6 plays a pivotal role in shaping the α-CTLA-4/α-PD-1-induced immunological landscape of rejecting T3 tumors. To better define the role of IP-CN6 in the T cell-dependent anti-tumor response, we clustered CD4^+^ and CD8^+^ T cells as Ki67^+^ (as a marker for T cell proliferation) versus GZMB^+^ (as a marker for cytotoxicity) **(Figure 6A)** and calculated their distribution among the seven CNs at the d10 time point **(Figure 6B and 6C)**. In control mice, Ki67^+^ and GZMB^+^ T cell populations were distributed in similar proportions at the tumor boundary (CN1) and within IP-CN6. However, in α-CTLA-4/α-PD-1-treated mice, both Ki67^+^ and GZMB^+^ CD4^+^ and CD8^+^ T cells sharply increased in frequency in IP-CN6 **(Figure 6B and C)**. This observation indicates that IP-CN6 is the preferential neighborhood where T-cell expansion and effector functions are facilitated in T3 tumors.

**Figure 6.**
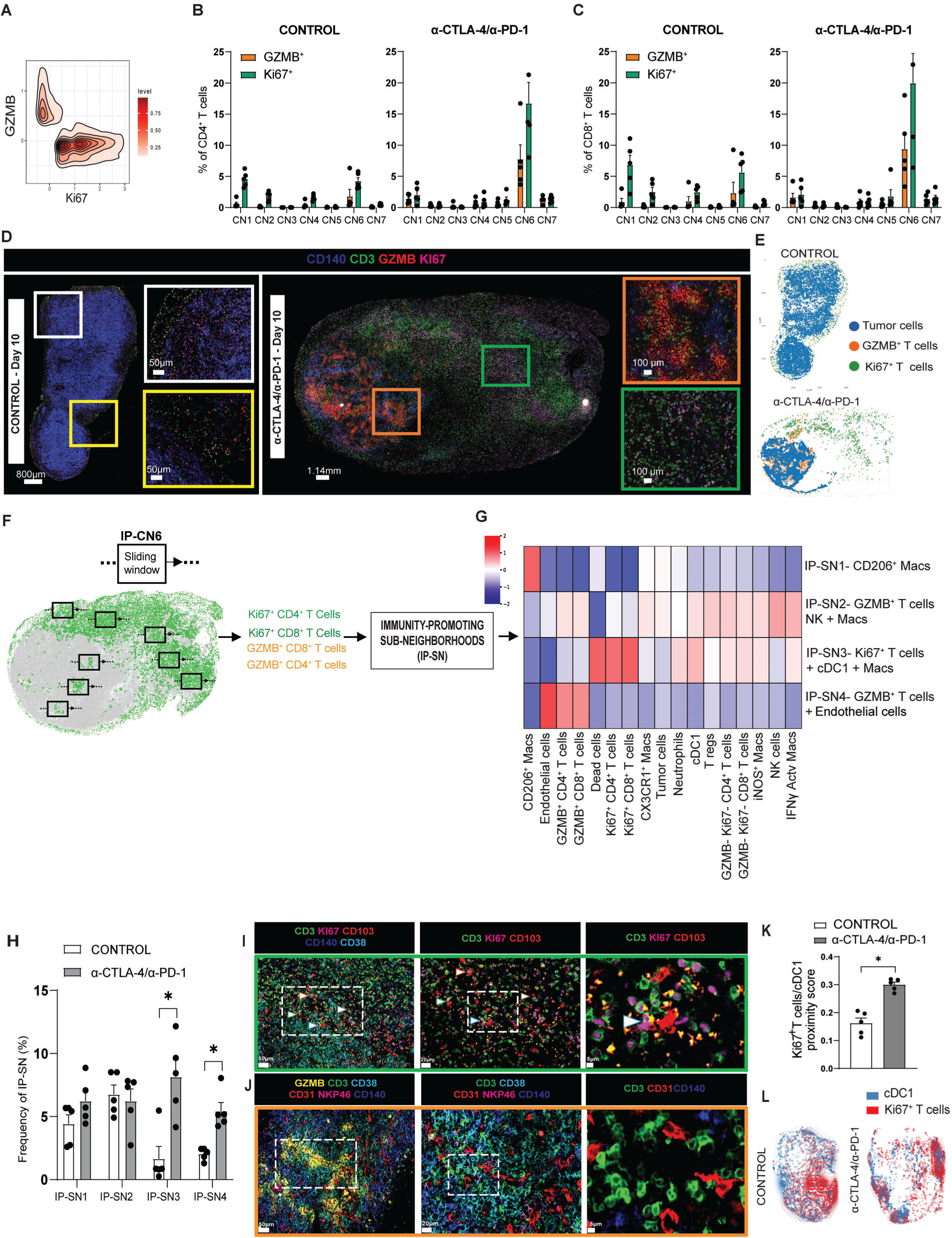
Granzyme B*⁺* and Ki67*⁺* T Cells Form Distinct Sub-Neighborhoods (IP-SNs) Within Immunity-Promoting CN6. **(A)** Representative contour plot showing GZMB^+^ versus Ki67^+^ T cells. **(B)** Frequency of GZMB^+^ and Ki67^+^ CD4^+^ T cells across the seven CNs in d10 T3 tumors from control (n=5) and α-CTLA-4/αPD-1 (n=5) mice. (**C)** Frequency of GZMB^+^ and Ki67^+^ CD8^+^ T cells across the seven CNs in d10 T3 tumors from control (n=5) and α-CTLA-4/αPD-1 (n=5) mice. **(D)** Four-color overlay CODEX image displaying the location of CD140a^+^, T cells (CD3^+^), GZMB^+,^ and Ki67^+^ cells in T3 tumors harvested on d10. Scale bar as indicated. **(E)** Dot plot showing the location of tumor cells, GZMB^+^ T cells, and Ki67^+^ T cells in a T3 tumor harvested on d10. **(F)** Computational method used to identify the immunity-promoting sub-neighborhoods (IP-SNs). **(G)** Heat map showing the identification of the IP-SNs and their respective frequencies (enrichment score) within each IP-SN (pooled data from both groups). **(H)** Frequency of the four IP-SNs in T3 tumors from control (n=5) and α-CTLA-4/αPD-1 (n=5) treated mice. *p < 0.05, Multiple unpaired T-test. (**I)** Green box showing a five-color CODEX image showing interactions between Ki67^+^T cells and cDC1(CD103^+^) (highlighted in white arrows with colored outlines) with a sequential increase in magnification. Scale bar goes from 50μm, 20μm and 5μm (left to right). **(J)** Orange**-**box showing six-color overlay CODEX image with a sequential increase in magnification for the cells in the vicinity of GZMB^+^ T cells. Scale bar goes from 50μm, 20μm and 5μm (left to right). **(K)** Proximity density analysis between Ki67^+^ T cells and cDC1. *p<0.05, T-test. **(L)** Contour plot showing the distribution of Ki67^+^ T cells (red) and cDC1 (blue) in a T3 tumor harvested on d10.

However, upon more detailed inspection, we detected differential spatial patterns between Ki67^+^ and GZMB^+^ CD3^+^ T cells between control mAb versus α-CTLA-4/α-PD-1 mice **(Figures 6D and 6E)**. In tumors from control mAb-treated mice, both populations were intermixed and resided at the outer region (periphery) of the tumor tissue. In contrast, following α-CTLA-4/α-PD-1, Ki67^+^ T cells remained concentrated at the tumor periphery while GZMB^+^ T cells infiltrated into the tumor core.

This observation suggests a more granular spatial organization within IP-CN6. To investigate this, we re-annotated both CD4^+^ and CD8^+^ T cells as either double-negative or single-positive for Ki67 or GZMB, performed sliding window mapping analysis exclusively within IP-CN6, and identified structures that we termed immunity-promoting sub-neighborhoods (IP-SNs) **(Figure 6F)**. This analysis revealed that IP-CN6 is comprised of 4 distinct IP-SNs **(Figure 6G)**. IP-SN1 contained CD206^+^ macrophages but did not show a significant change in tumors from control versus α-CTLA-4/α-PD-1 mice **(Figure 6H)**. IP-SN2 was composed of GZMB^+^ CD4^+^ T cells, GZMB^+^CD8^+^ T cells, NK cells, and IFNγ-activated macrophages **(Figure 6H),** but also did not significantly change following α-CTLA-4/α-PD-1 versus control mAb therapy. However, IP-SN3 contained a significant enrichment of proliferating T cells, IFNγ-activated macrophages and cDC1 following α-CTLA-4/α-PD-1 compared to control Ab. IP-SN4 showed significant α-CTLA-4/α-PD-1-induced increases in cytotoxic GZMB^+^ T cells and endothelial cells compared to control **(Figure 6H).**

Inspection of d10 tumors from α-CTLA-4/α-PD-1 mice reveals the differences in cellular composition surrounding Ki67^+^ versus GZMB^+^ T cells. Ki67^+^ T cells were predominantly found at the periphery of the tumor, were in proximity to IFNγ-activated macrophages and were observed interacting with cDC1s (as indicated by the white arrows). **(Figure 6I)**. The co-localization between cDC1 and Ki67^+^ T cells was additionally evidenced by an increased proximity score from 0.15 (control) to 0.3 (α-CTLA-4/α-PD-1) **(Figure 6K and 6L**). In contrast, GZMB^+^ T cells were found infiltrating the tumor core and forming an intratumoral niche with IFNγ-activated macrophages, NKP46^+^ NK cells, and CD31^+^ endothelial cells. **(Figure 6J).** Overall, these findings support our hypothesis that IP-CN6 constitutes a specialized microenvironment for myeloid APC and T cells that facilitates the acquisition of proliferative versus cytotoxic functions following α-CTLA-4/α-PD-1. However, the spatial compartmentalization and formation of sub-neighborhoods suggest that the induction of these functions during tumor rejection follows distinct dynamics.

### Temporal Remodeling of IP-CN6 Drives the Spatial Compartmentalization Between Cytotoxic and Proliferative Functions During *α*-CTLA-4/*α*-PD-1

To resolve the question of how the kinetics of compartmentalization of Ki67^+^ T versus GZMB^+^ cells evolved over time, we analyzed the dynamics of cytotoxicity and proliferation of CD4^+^ and CD8^+^ T cells within IP-CN6. In tumors from control mice, T cells gradually lost expression of Ki67 and GZMB after d9 **(Figure 7A and 7B)**. However, after α-CTLA-4/α-PD-1, a surge in Ki67^+^ CD4^+^ and CD8^+^ T cells was observed on d10, followed by an increase in GZMB expression on d11 and d13 **(Figure 7A and 7B)**. This indicates that T cells first undergo a proliferative burst before expressing increased cytotoxic activity on d11, leading to complete tumor elimination by d13.

**Figure 7.**
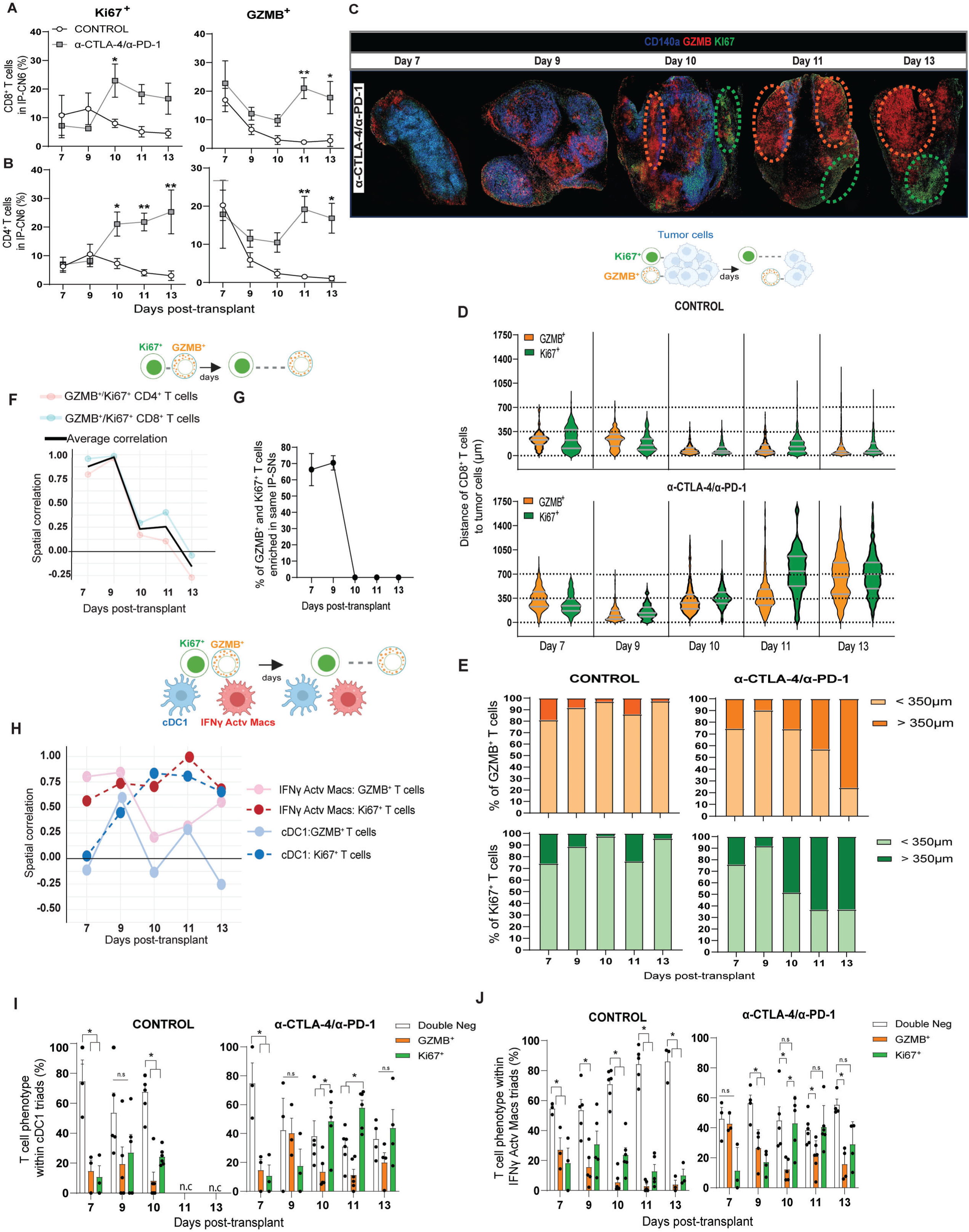
Temporal Remodeling of IP-CN6 Drives the Spatial Compartmentalization Between Cytotoxic and Proliferative Functions During *α*-CTLA-4/*α*-PD-1. **(A)** Longitudinal frequency of GZMB^+^ and Ki67^+^ CD8^+^ T cells in the IP-CN6. *p<0.05, **p=0.0094. ANOVA (with mixed-effects analysis) **(B)** Longitudinal frequency of GZMB^+^ and Ki67^+^ CD4^+^ T cells in the IP-CN6. *p<0.05, **p=0.0099. ANOVA (with mixed-effects analysis). From A to B, mean in Control represents [n=3(d7), n=4(d9), n=6(d10), n=5(d11), n=3(d13)] and in α-CTLA-4/αPD-1 [n=3(d7), n=4(d9), n=5(d10), n=5(d11), n=4(d13)]. **(C)** Longitudinal three-color CODEX images showing the distinct location of GZMB^+^ and Ki67^+^ cells in T3 tumors from α-CTLA-4/α-PD-1 mice. **(D)** Violin plots depicting the distance of GZMB^+^ or Ki67^+^ CD8 T cells to tumor bulk cells over multiple time points. **(E)** Percentage of GZMB^+^ or Ki67^+^ CD8^+^ T cells above or below 350 μm to tumor bulk cells. **(F)** Spatial correlation between the location of Ki67^+^ and GZMB^+^ T cells inside the IP-CN6 over multiple time points. A high spatial correlation indicates that these cell types are located near one another (i.e., within the same IP-SNs), while a low or negative correlation suggests they are distributed across distinct IP-SNs and spatially distant. **(G)** % of GZMB^+^ or Ki67^+^ CD8 T cells enriched within the same IP-SNs over time. **(H)** Spatial correlation between the location of cDC1 and IFNγ-activated with respect to Ki67^+^ or GZMB^+^ T cells inside the IP-CN6 over multiple time points. **(I)** T cell phenotype in the vicinity of cDC1 triads. n.c= not calculated *p<0.05, Tukey’s multiple comparisons test **(J)** T cell phenotype in the vicinity of IFNγ-activated triads. *p<0.05, Tukey’s multiple comparisons test

To further understand the dynamics of induction of effector function, we analyzed the expression levels of PD-1, LAG-3, and TIM-3 on CD4^+^ and CD8^+^ T cells to define their activation versus exhaustion states **(Figure S11A)**. No significant differences in single expression of PD-1 were observed on either CD8^+^ and CD4^+^ T cells between tumors from control mAb and α-CTLA-4/α-PD-1 mice **(Figure S11B and S11C**, respectively). However, after d10, we noted a statistically significant increase in terminally exhausted CD8^+^ T cells (PD1^+^LAG3^+^ and PD-1^+^LAG-3^+^TIM-3^+^) in tumors from control mAb-treated mice, but not in tumors from α-CTLA-4/α-PD-1-treated mice **(Figure S11D and S11F**, respectively). We did not see significant changes in terminal exhaustion markers in CD4^+^ T cells in tumors from either control mAb or α-CTLA-4/α-PD-1-treated mice **(Figure S11E and S11G)**. These results indicate that the continuous increase in T cell effector function seen in IP-CN6 following α-CTLA-4/α-PD-1 therapy is accompanied by reduced cellular exhaustion. The differences observed between CD8^+^ and CD4^+^ tumor-associated T cells were most likely attributable to the fact that in T3 tumors, 40% of CD8^+^ T cells recognize the MHC-I-restricted neoantigens mLama4 and mAlg8, while only 6% of CD4^+^ T cells recognize the major MHC-II neoantigen, mItgb1^10, 13^.

Longitudinal analysis of Ki67^+^ and GZMB^+^ T cells suggested that their distinct spatial localization had not occurred at d7 and d9, but became detectable at d10, (at a time when tumor cell killing first became obvious) and then was maintained on d11 and d13 **(Figure 7C)**. To assess when GZMB⁺ and Ki67⁺ T cells are found in different regions of the tumor tissue, we first calculated the full distance distributions of effector and proliferating T cell populations to bulk tumor cells over time **(Figure 7D).** We then summarized the fractions of Ki67^+^ or GZMB^+^ T cells located within a mid-distribution reference point, of close range from tumor cells (below 350μm) or distant from tumor cells (above 350μm) **(Figure 7E).** On d9, approximately 80% of Ki67^+^ or GZMB^+^ CD8^+^ T cells in tumors from either control mAb or α-CTLA-4/α-PD-1 mice were located within 350μm of tumor cells. However, on d10, while GZMB^+^ cells remained in proximity to tumor cells, 50% of Ki67^+^ T cells in tumors from α-CTLA-4/α-PD-1 mice were found at the tumor periphery at a distance greater than 350μm from tumor cells **(Figure 7E)**. This spatial compartmentalization continued on d11, with 70% of Ki67^+^ cells accumulating further away from bulk tumor cells, whereas 60% of GZMB^+^ cells remained within 350μm of their targets. It was only on d13, when tumor cell elimination was nearly complete, that GZMB^+^ T cells dispersed and were found at distances greater than 350μm away from their tumor target cells **(Figure 7E)**.

For the second approach, we employed the sliding window mapping method exclusively within IP-CN6 at individual time points after α-CTLA-4/α-PD-1 **(Figure S12A)**. This allowed us to identify IP-SNs on specific days but not shared across all time points. On d7 and d9, the IP-CN6 contained two IP-SNs where both Ki67^+^ and GZMB^+^ T cells co-localized into the same cluster **(Figure S12B and S12C**, respectively). However, d11 and d13 after α-CTLA-4/α-PD-1, two IP-SNs evolved, showing selective spatial segregation of Ki67^+^ versus GZMB^+^ T cells **(Figure S12D and S12E,** respectively). To better define this observation, we quantified the spatial correlation between the location of proliferating versus effector T cells within the IP-SNs after α-CTLA-4/α-PD-1 over time. In this context, a high spatial correlation indicates that these cell types are located near one another (i.e., within the same IP-SNs), while a low or negative correlation suggests they are distributed across distinct IP-SNs and spatially distant. Consistent with our observations, Ki67⁺ and GZMB⁺ T cells exhibited strong spatial overlaps early after treatment, which sharply declined after d10 and reached a minimum by d13 **(Figure 7F)**. This dynamic was evidenced by the proportions of T cells co-enriched within the same IP-SNs of IP-CN6: on d7, where approximately 66% of Ki67⁺ and GZMB⁺ T cells were found within the same IP-SNs, increasing to 77% by d9, and then rapidly declining after d10 **(Figure 7G)**. These findings demonstrate a dynamic transition from overlapping proliferative and cytotoxic compartments to spatially segregated functional niches.

To determine how myeloid populations contribute to this spatial compartmentalization, we examined the spatial relationships between cDC1s, IFNγ–activated macrophages with Ki67⁺ and GZMB⁺ T cells **(Figure 7H)**. On d9, both myeloid APCs were spatially associated with proliferating and cytotoxic T cells. However, after d10, this spatial association was only maintained for Ki67⁺ T cells, while the correlation with GZMB⁺ T cells declined. To further define this association, we characterized the T cell phenotype in the vicinity of cDC1 and IFNγ–activated macrophages engaged in triad interactions (previously identified in **5L and 5M**). In control tumors, cDC1 triads were primarily associated with non-proliferating, non-cytotoxic T cells as shown on d10 (**Figure 7I)**. Later timepoints were not analyzed, as triad-forming cDC1s were no longer detected beyond this stage. Following α-CTLA-4/α-PD-1, however, cDC1 triads became preferentially associated with Ki67⁺ T cells, particularly on d10 and d11 (**Figure 7I)**. IFNγ–activated macrophage triads followed a similar pattern. In tumors from control mice, they were predominantly associated with double-negative cells, but after α-CTLA-4/α-PD-1, the proportion of Ki67⁺ T cells within these triads increased markedly, especially on d10 and d11, and exceeded the frequency of GZMB⁺ T cells (**Figure 7J)**. These findings suggest that cDC1s and IFNγ–activated macrophages within IP-CN6 establish a specialized sub-neighborhood that supports intratumoral T cell proliferation.

Taken together, these results reveal a dynamic maturation process within IP-CN6 that underlies effective tumor rejection. Early after T cell infiltration (d7–d9), proliferating and cytotoxic T cells co-localize within shared sub-neighborhoods. As tumor elimination begins, these populations become spatially segregated, with proliferating T cells remaining in APC-triad-rich niches at the tumor periphery, while CD8⁺ T cells target residual neoplastic cells and maintain cytotoxic activity in the proximity of tumor cells. This spatial and functional compartmentalization was not observed in progressively growing tumors, indicating that the emergence of APC-supported proliferative niches and organized effector activity is a defining feature of successful ICT.

### The Maturation of the Immunity-Promoting CN6 Reshapes Tissue Organization and the Communication Rules Between CNs

Whereas the previous findings revealed internal T-cell spatial dynamics occurring during the maturation of the IP-CN6, it remained unclear how the increase of the IP-CN6 impacts tissue organization and how it interacts with the remaining CNs, particularly with the tumor boundary and tumor bulk CNs, which are enriched for tumor cells. To answer this question, we first sought to define the communication rules between the seven CNs and tested the hypothesis that CN interactions were more likely to happen in regions where their cellular members are physically closer to the interfaces between adjacent CNs **(Figure 8A)**. These interfaces are thought to be sites where molecules produced by one CN can signal to another CN, thus enabling inter-CN communication^36^. To identify cells at the interfaces of adjacent CNs rather than their own CN, we used a 20-nearest-neighbor analysis to identify cells with at least 30% of their neighbors belonging to an adjacent CN as interface cells. We then used a permutation test to shuffle the interface cells and generated a communication network model, where each node represents a CN, and arrows represent statistically enriched adjacent at CN interfaces. Importantly, these interactions reflect spatial opportunities for communication, rather than measured ligand–receptor engagement **(Figure 8A)**.

**Figure 8.**
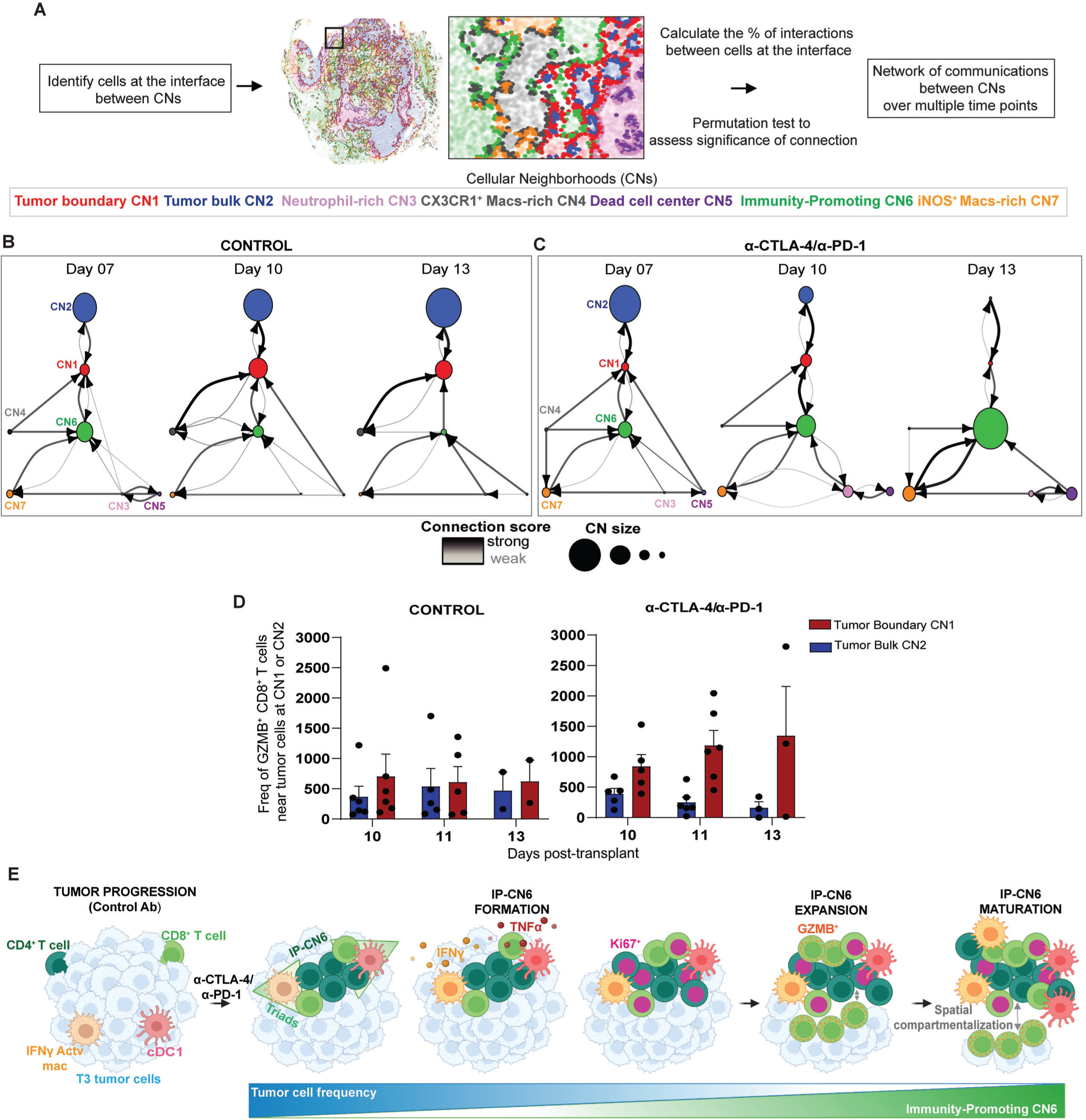
The Maturation of the Immunity-Promoting CN6 Reshapes Tissue Organization and the Communication Rules Between CNs. **(A)** Computational method utilized to identify cells at the interface of CNs and build a network of communication between CNs. Node size reflects CN frequency in the tissue, and connection lines indicate connection strength, ranging from weak (light gray) to strong (black)**. (B)** Network of communication between the CNs in T3 tumor from control Ab mice. **(C)** Network of communication between the CNs in T3 tumor from α-CTLA-4/α-PD-1 mice. **(D)** Frequency of GZM-B^+^ CD8 T cells to tumor cells at CN1 and CN2. **(E)** Proposed spatiotemporal mechanism for the maturation of IP-CN6 during α-CTLA-4/α-PD-1.

In tumors from control mice **(Figure 8B)**, tumor bulk CN2 cells exclusively communicated with tumor boundary CN1 cells, positioning the boundary CN as a node connecting both the CX3CR1^+^ macrophage-rich CN4 and IP-CN6. On d7, the CX3CR1^+^ macrophage-rich CN4 was a minor component, but by d13, it became the primary node interacting with the tumor boundary CN1, while communication with IP-CN6 diminished concomitantly. Following α-CTLA-4/α-PD-1 **(Figure 8C)**, the order of interactions between tumor bulk CN2 and boundary CN1 remained unchanged. However, as IP-CN6 expanded from d7 to d13, it assumed a central role, becoming the strongest communication node with the tumor boundary CN1 while also communicating with nearby myeloid CNs, including the iNOS^+^ macrophages-rich CN7 and Neutrophil-rich CN3. Notably, the latter two CNs were not shown to be communicating with the tumor boundary, especially in tumors from control mAb-treated mice, which were characterized as having a small node size, reflecting the low frequency of these CNs.

These findings suggest preferential communication patterns between different CNs but do not identify which CN proximity relationships were most affected by α-CTLA-4/α-PD-1. We therefore analyzed the collective behavior of the cell types comprising each CN. Cells were labeled according to their respective CNs, 20-nearest-neighbor analysis was performed across multiple time points (individually represented in **Figure S13)** and a statistical analysis of all CN nearest-neighbor interactions across all time points (summarized in **Figure S14)**. In tumors from mice treated with control mAb, the top three interactions were (highlighted in the green box at the bottom of the heatmap): iNOS^+^ macrophages-rich CN7 and CX3CR1^+^ macrophage-rich CN4, (ii) tumor boundary CN1 and CX3CR1^+^ macrophage-rich CN4, and (iii) IP-CN6 and CX3CR1^+^ macrophage-rich CN4. In contrast, the top three interactions following α-CTLA-4/α-PD-1 were (highlighted in purple box at the top of the heatmap): (i) IP-CN6 interacting with tumor boundary (CN1), (ii) IP-CN6 interacting with the iNOS^+^ macrophages-rich CN7, and (iii) IP-CN6 interacting with CX3CR1^+^ macrophage-rich CN4.

Collectively, these results define the communication rules governing inter-CN interactions and reveal a spatial hierarchy that emerges during tumor rejection. In control mAb-treated tumors, the tumor boundary CN1 was progressively associated with the CX3CR1⁺ macrophage–rich CN4, which competed with IP-CN6 for connection to the tumor boundary. In contrast, following α-CTLA-4/α-PD-1 therapy, IP-CN6 became the dominant communication node in the tissue, preferentially interacting with the tumor boundary CN1 rather than the tumor bulk CN2, while also maintaining connections with myeloid CN4 and the iNOS⁺ macrophage–rich CN7.

To determine whether this IP-CN6 preferential communication with CN1 resulted in localized effector function, we performed nearest-neighbor analysis on tumor cells located at CN1 versus CN2 and quantified the frequency of GZMB⁺ CD8⁺ T cells in their proximity over time **(Figure 8D)**. This analysis revealed a significant increase in the frequency of cytotoxic CD8⁺ T cells only after α-CTLA-4/α-PD-1 and specifically in proximity to the tumor cells at the tumor boundary CN1, with no corresponding increase in the tumor bulk CN2. These findings suggest a sequential mechanism of tumor rejection in which effector CD8⁺ T cells from IP-CN6 preferentially target the tumor boundary CN1, where they can progressively eliminate layers of tumor cells as the IP-CN6 expands during α-CTLA-4/α-PD-1 therapy and reorganizes the TME.

## DISCUSSION

In this study, we applied longitudinal spatial proteomic profiling to define the immune dynamics underlying tumor rejection following dual α-CTLA-4/α-PD-1 immunotherapy. By preserving tissue architecture, this approach extends our prior CyTOF and scRNA-seq analyses of disaggregated tumors^14^. Using the T3 tumor model with well-defined rejection kinetics, we analyzed multiple cohorts across different time points, enabling comprehensive characterization of the rejection process and the development of a spatiotemporal mechanistic model for the evolving role of an intratumoral CN composed of CD4⁺ and CD8⁺ T cells engaging in triad interactions with cDC1s or IFNγ–activated macrophages **(Figure 8E)**. In this model, progressively growing tumors have T cells that are scarce, poorly infiltrate tumor tissue, and rarely engage cDC1s or IFNγ–activated macrophages, resulting in minimal formation of APC-CD4-CD8 cellular triads. Following α-CTLA-4/α-PD-1 therapy, however, the TME undergoes coordinated reorganization, enabling the dynamic evolution of IP-CN6, which progresses through three functionally distinct phases. During the **formation phase**, CD4⁺ T cells first increase in frequency and spatially redistribute, enabling co-localization with CD8⁺ T cells and APCs and initiating assembly of IP-CN6. At this stage, T cells exhibit a burst of proliferation followed by early strong induction of TNFα and IFNγ, consistent with reciprocal activation of cDC1s and macrophages. During the **expansion phase**, CD8⁺ T cells markedly increase and surpass CD4⁺ T cells in frequency, coinciding with a peak in T cell cytotoxicity and substantial tumor cell elimination. This step is accompanied by continued T cell and APC engagement and an increase in triad frequency. Finally, during the **maturation phase**, spatial compartmentalization becomes evident: Ki67⁺ proliferating T cells localize preferentially near triad-forming APCs—often at the tumor periphery—while GZMB⁺ cytotoxic T cells move inward from the tumor boundary CN1 and target remaining tumor cells. Concomitantly, this transition is associated with a reorganization of inter-neighborhood interactions, establishing IP-CN6 as a dominant structural and functional hub within the tissue.

Our findings extend the prevailing model of tumor rejection, from the previous model involving a simple, quantitative increase in cytotoxic activity to a complex process in which the IP-CN6 functions as a lymphoid-like organ that dynamically organizes antitumor immunity. Moreover, our findings provide critical temporal resolution to our understanding of the role of multicellular neighborhoods and directly demonstrate how immune organization changes over time, determining tumor fate. Our study is among the first to apply spatial proteomic analysis across the entire span of the antitumor immune response, from initiation to complete elimination. In contrast, prior studies have largely relied on single-time-point analyses, providing static snapshots that cannot resolve when key cellular events take place^15, 37^. Together, our findings establish maturation of immune neighborhoods and functional compartmentalization as novel principles underlying successful ICT.

Mechanistically, our results indicate that the efficacy of α-CTLA-4/α-PD-1 relies on coordinated crosstalk between T cells and diverse APCs. The formation of two distinct triad types, prior to the expansion of CD8⁺ T cells, indicates that immune triads may be involved with the rapid proliferation and increase in cytotoxicity observed from d11 to d13. We hypothesize that initial tumor cell killing generates antigen release that is captured by APCs, enabling further CD8⁺ T cell expansion. This model aligns with established roles of cDC1s in cross-presentation and MHC-II–dependent CD4⁺ T cell help³⁸. Triads function by promoting physical proximity of T cells and APCs, facilitating chemokine and/or cytokine production and their biologic activities^31^. TNFα signaling via TNFR1 has been shown to induce cDC-1 maturation, as well as to enhance co-stimulatory activity³⁹,⁴⁰. TNFα has also been shown to enable cDC1s licensing of CD8^+^ T cell cytotoxicity³⁹. In addition, IFNγ-dependent modulation/activation of myeloid cell function is well known^38^. IFNγ-polarized macrophages have been shown to directly kill certain tumor cells, enhance T cell cytotoxicity by functioning as APCs and secreting pro-inflammatory cytokines⁴¹,⁴². These mechanisms would provide a positive feedback loop in which activated T cells and myeloid APCs reinforce each other’s functions to sustain a productive immune response.

However, not all macrophage subsets contribute positively. CX3CR1⁺ macrophages in tumors from control mAb-treated mice represent immature myeloid populations with limited co-stimulatory and antigen-presenting capacity, as shown by others^39, 40, 41^. Similarly, CD206⁺ macrophages are often considered to express immunosuppressive functional activities^42^. Yet, emerging evidence indicates that CD206⁺ macrophages can efficiently capture antigens and stimulate antigen-specific CD8⁺ T cells^43^. This would be consistent with our observation that CD206⁺ macrophages are present in IP-CN6 and may support T cell activity, though their function remains unclear in our model.

While organized networks of T cells and APCs have been previously described in other mouse models, their temporal evolution has not been demonstrated. In the Kras/Trp53 (KP) lung adenocarcinoma model, T and B cells form organized structures in response to immunogenic epitopes, yet these tumors remain refractory to ICT^22^. Similarly, Hickey et al. observed in a model of adoptive T cell therapy that organized CD8⁺ T cell zones drove upregulation of PDL1 on macrophages, reduced tumor proliferation, but ultimately failed to eliminate tumors^44^. By contrast, we observed that following α-CTLA-4/α-PD-1, IP-CN6 drives rapid and complete tumor rejection, without enforced antigen expression or requiring B cells, as we previously showed that T3 tumors do not elicit a B cell response^14^.

Analogous to these observations, in human cancers, Chen et al. identified “immunity hubs” enriched in stem-like CD8⁺ T cells and activated DCs in lung cancer, correlating with responses to anti-PD-1^23^. However, their analyses were limited to pre-treatment samples, leaving unknown the temporal trajectory of these structures. Shiao et al. identified an ‘immune district’ in breast cancer patients that expanded in response to pembrolizumab and radiation therapy, characterized by effector T cells, B cells, and MHCII^+^ macrophages^17^. Yet, the single-arm design makes it difficult to differentiate anti-PD-1’s specific effect.

In conclusion, this study reveals a spatiotemporal mechanism that shows how a neighborhood of T cells and myeloid APCs emerges, expands, and functionally evolves in response to α-CTLA-4/α-PD-1. Our longitudinal CODEX dataset highlights the power of spatial proteomics to identify dynamic immune organization and provides a valuable resource to obtain mechanistic insights. The current study will thus provide a critical dataset to compare the effects of mono-ICT to dual ICT, and between ICT and cancer vaccine therapy. Future spatial transcriptomic analyses will also resolve how the two distinct triad configurations contribute to CD8⁺ T cell programming and will aim to generate a multi-omics approach to refine our observations. A particularly important focus of our future research is to explore whether neighborhoods similar to IP-CN6 arise in additional tumor models across diverse mouse strains, including those in which MHC-II is expressed directly by the tumor cell and whether such structures can be induced by other immunotherapies^12, 45, 46^. Defining which tumor contexts and therapeutic strategies elicit these immune architectures will be critical for developing the next generation of cancer immunotherapies.

## Acknowledgments

We thank Yury Goltsev for providing technical and scientific assistance. This work was supported by grants to R.D.S. from the National Cancer Institute of the US National Institutes of Health (R01CA190700), the Parker Institute for Cancer Immunotherapy, and a Stand Up to Cancer-Lustgarten Foundation Pancreatic Cancer Foundation Convergence Dream Team Translational Research Grant. Stand Up to Cancer is a program of the Entertainment Industry Foundation administered by the American Association for Cancer Research. The IML is supported by the Andrew M. and Jane N. Bursky Center for Human Immunology and Immunotherapy Programs and the Alvin J. Siteman Comprehensive Cancer Center. The latter is supported by a National Cancer Institute of the National Institutes of Health Cancer Center support grant (P30CA91842).

## Author contributions

R.F.V.M. and R.D.S. conceived and designed the experiments, interpreted the results, and wrote the manuscript. R.F.V.M. developed the Ab panel, performed the experiments, collected and analyzed the data. R.D.S. oversaw all aspects of the study. M.N.A. oversaw the bioinformatics analysis and interpreted the results. V.S. performed bioinformatics analysis and interpreted the results. K.H-H and B.R performed ARCADIA and ContactTracing analysis and discussed the data. F.H. assisted with experiments and ab panel development. B.B.C. developed the singer imaging processing software and provided technical assistance. H.S. performed the neutrophil depletion experiment. H.S and Y.S. helped with the scRNA-seq experiment. A.S. provided advice on analytical aspects of the project. B.H.Z. provided critical imaging assistance. D.A., S.A., D.J.T., and Y.T assisted with sample collection and provided technical assistance. K.C.F.S. provided critical review of the manuscript. M.W., and H.K. managed the lab and oversaw the mouse colony. P.L.jr, and P.Lsr. provided the CODEX instrument and custom-made CODEX antibodies. G.N. provided methodology and analysis assistance. E.A. supervised ARCADIA and ContactTracing analysis, provided analysis assistance, and discussed the data. All authors contributed to the final development of the manuscript and agreed with the results and conclusions.

## METHODS

### Mice

Six-week-old wild-type 129S6 male mice were purchased from Taconic Farms. *Rag2^−/−^* male 129S6 mice were bred in our specific-pathogen-free facility. Experiments were done on 8-week-old males. All studies were performed in accordance with procedures by AAALAC accredited Animal Studies Committee of Washington University in St. Louis.

### Tumor transplantation

In this study, we used the T3 MCA-induced sarcoma line, derived from a male 129S6 mouse which was previously characterized in^10^. Tumor cell aliquots were kept in frozen stocks and cultured *in vitro* with RPMI media (Hyclone) supplemented with 10% FCS (Hyclone). For tumor inoculation, T3 tumor cells were resuspended at 1.0 ×10^7^ cells per mL in endotoxin-free PBS and then 1.5 × 10^6^ cells were injected subcutaneously into the flanks of mice (150μl). Viability was assessed with AOPI before injection and tumor cells with >90% viability were injected into mice. Tumors were measured every 3 days by caliper measurements and calculated as the average of two perpendicular diameters.

### Immune checkpoint therapy (ICT)

T3 tumor-bearing mice were injected intraperitoneally with the combination of 200 μg of α-CTLA-4 (9D9) and 200 μg α-PD-1 (RMP1-14), or 200 μg of rat IgG2a isotype control mAb as control, on days +6, +9, +12 post-tumor transplant. Endotoxin-free antibodies were diluted in PBS before injection to a total volume of 0.5 mL per mouse.

### Tumor harvest for flow cytometry

On day +10 post-tumor transplant, established T3 tumors were excised from mice, chopped, and treated with 1 mg/ml type 1A collagenase (Sigma) in HBSS (Hyclone) for 1 hour at 37 °C as previously described^14^. For conventional flow cytometry staining, single cell suspensions of digested tumors were stained for 5 minutes at room temperature with 500 ng of FC block (anti-CD16/32) in 100 μL of staining buffer (PBS with 2% FCS) and then stained with surface antibodies for 30 minutes at 4 °C as described in STAR methods for antibodies concentration. Flow cytometry was performed on the LSR Fortessa X-20 (BD Biosciences) and analyzed with the FlowJo software (Treestar).

### Fresh frozen tumors

On days +7, +9, +10, +11, and +13 post-tumor transplant, established T3 tumors were excised from mice, placed on a labeled standard Tissue-Tek Cryomold (25mm x 20mm x 5mm, Sakura cat# 4557), and covered with Tissue-Tek O.C.T. compound (Sakura cat#4583) for 1-2 minute at room temperature to acclimate the sample before being transferred to the top of a cold metal block inside a bucket filled with enough liquid nitrogen to make contact with the middle/bottom part of the metal block. After that, the bucket was kept closed, and samples were left on top of the metal block for about 5 minutes to snap-freeze. Frozen samples were then rapidly wrapped with a pre-labeled aluminum foil, transferred to a foam cooler filled with dry ice, and stored at –80 °C freezer.

### Poly-lysine coverslip preparation

Coverslips were prepared according to the CODEX user’s manual. Briefly, glass cover coverslips (thickness 1 ½, 22 x 22 mm, Corning Cat #2850-22) were placed at the bottom of a glass beaker (0.5 L), dispersed by rotating the beaker, and covered with 20 mL of poly-lysine solution at RT for a minimum of 12 hours. After incubation, the poly-lysine solution was removed from the beaker, and the coverslips were washed with doublely distilled water (ddH_2_0) for a total of 7 washes (1 minute each). After this, coverslips were placed on top of a sheet of kimwipes, dispersed to avoid overlapping and left to dry before being transferred to a petri dish for storage

### Tissue cryosectioning

Tissue cryosectioning was done in a Leica Cryostat (CM 1860) at −20 °C. Briefly, frozen tissue blocks were transferred from the −80 °C freezer, left inside the Cryostat for 30 minutes to acclimate, and then cut into 8 μm thick sections. Sections were maneuvered on top of the coverslip and kept flat. The tissue was attached to the coverslip, transferred to a pre-labeled box, and stored at −80 °C. Alternatively, tissue sections were also transferred to Superfrost Plus Gold Microscope Slides (25 x 75 x1.0mm, Fisher Brand, Cat #15-188-48) to perform immunofluorescence staining.

### CODEX staining

CODEX staining was done according to the user’s manual for fresh frozen tissue sections. Coverslips kept at −80 °C freezer were transferred in a 50 mL Falcon tube filled with fresh drierite absorbents (Fisher Scientific, cat #23-116582) into a 50 mL glass beaker containing 10 mL of acetone (Sigma, cat # 472301-4L) and incubated for 10min at room temp. After that, tissue was removed from acetone, left to dry for 2 min and then transferred into a 6-well TC plate dish containing Hydration Buffer (∼5 mL/well). Coverslips were incubated for 2 min, transferred to another well containing Hydration Buffer (∼5 mL/well) for another 2 min, and then placed for 10 min in a new well-containing 5mL of pre-staining fixing solution which was 1 part 16% PFA (Electron Microscopy Sciences, Cat # 15710-S) in 9 parts of Hydration buffer at 1:10 (v/v). Next, the coverslips were washed two times with Hydration buffer and transferred to a new well containing Staining buffer to equilibrate for 30 min. During this time, the CODEX blocking buffer solution was prepared by mixing Staining Buffer (190 μL), N Blocker (5 μL), G blocker (5 μL), J Blocker (5 uL), S blocker (5 μL) and then by removing the total antibody volume (sum of each antibody volume) from the 210 μL CODEX blocking buffer final volume. CODEX antibodies are added according to the STAR methods dilution factor to make the 210 μL antibody cocktail solution, which was enough to stain 2 coverslips with 100 μL each using the Omni-Stainer™ C12 (Parhelia Biosciences cat # 40101). For this step, 100 μL of Staining solution was placed on top of each support tile and the tissue coverslip was carefully placed facing the support tile to create a capillarity chamber with 50 μm thin spacers. The support tile was then transferred to the built-in wet chamber and 100 μL of the Antibody cocktail solution was dispensed at the top edge of the coverslip to instantly wash out the staining solution by the laminar flow and form a capillary layer in which the sample was incubated overnight at 4°C. Coverslips were then removed from the support tile, washed two times in a 6-well plate containing staining buffer (5 ml), and fixed for 10 min with post-staining fixing solution (5 mL of 1 part 16% PFA in 9 parts of Storage buffer at 1:10 (v/v)). Coverslips were then quickly washed three times in 5 ml of 1x PBS, fixed with 5mL of cold Methanol (Honeywell cat#67-56-1) for 5 min while the 6-well plate sat on ice and then washed again three times in 5 mL of 1x PBS (Hyclone, cat# SH30028.03) before 20 min incubation with 200 μL (for each coverslip) of freshly made Final Fixative solution which is 1mL of 1X PBS and 20 μL of Fixative reagent (Akoya). Samples were washed subsequently three times in 5 mL of 1x PBS and stored for up to two weeks in 5 mL of storage buffer at 4°C.

### Reporter plate preparation

Reporter stock solution was made according to the CODEX user’s manual for 15 cycles which is 3660 mL of Nuclease-free water (Invitrogen™, cat # AM9930), 450 μL of 10X CODEX Buffer, 375 μL of Assay Reagent, and 15 μL of Nuclear Stain. After mixing, the reporter stock solution was transferred to labeled amber tubes, and 5 μL of each reporter dye corresponding to each imaging cycle was added to complete the final volume of 245 μL of the Reporter Master Mix and then transferred to 96-well plates.

### Generation of custom-conjugated CODEX DNA-conjugated antibodies

All Ab clones used for custom-conjugation came from commercially validated vendors and had their specificities confirmed by us through standard immunofluorescence staining before conjugation. If available, we conjugated the same ab clones previously used for flow cytometry or mass cytometry analyses^14^. Additionally, monoclonal Abs recommended for fresh immunofluorescence were used preferentially over polyclonal Abs (for details on clones and manufacturers, see **Table S1)**. To ensure high yields of conjugated antibodies, with no detectable free antibody or free barcode, all custom conjugations were performed by Leinco Technologies and carried out with 100 μg of carrier-free antibody using the custom-conjugation kit (Akoya Biosciences, Cat# PCSP0200). For optimal staining, we chose to conjugate antibodies targeting high-abundance proteins to Alexa 488-labeled DNA oligonucleotides. Antibodies targeting low-abundance proteins or nuclear markers were preferentially conjugated to DNA oligonucleotides labeled with ATTO550 or Alexa647. Custom and commercially available clones were validated by staining mouse spleen or tumor tissue. CD140a (PDGFR-α) was used as a marker for T3 tumor cells. PDGFR-α is highly expressed by T3 tumor cells with a 200.296 FPKM expression level (based on cDNA-Cap-seq data)^7^. Tumor cell staining with CD140a mAb was confirmed using OFP-expressing T3 tumor cells and observing a high degree of colocalization**(Figure S1B)**. Non-malignant PDGFR-α^low^ stromal cells were rare and showed discordant morphology and marker context. For the other markers in the CODEX Ab panel, cell morphology and co-staining patterns were compared with the literature and confirmed. The expected antigen co-staining patterns can be observed using control and α-CTLA-4/α-PD-1-treated samples **(Figure S3 A and B**).

### CODEX multiplex imaging

Imaging was done using a DMi8 Leica fluorescent microscope configured with three fluorescent channels (Alexa-488, ATTO 550, Cy5) and equipped with a 20x/0.80 air immersion apochromatic corrected objective with high numerical aperture and high resolution. Automated image acquisition, hybridization, and stripping of the fluorescent oligonucleotides were performed using a CODEX instrument (currently rebranded as Phenocycler®) and the CODEX Instrument Manager Software (Akoya Biosciences, version 1.30.0.12), which are commercialized by Akoya Biosciences (currently acquired by Quanterix). Imaging cycles were typically set to 17. The specific order of reporters, along with their corresponding exposure times, can be found in detail within **Table S2**. Additionally, we followed the recommendations described in Black et al^25^ for the optimal order of markers, particularly considering nuclear markers like Foxp3 and other low-expressing markers. Notably, these markers were preferentially imaged during the initial cycles of the process. To prepare for a CODEX run, bottle 1 of the CODEX instrument was filled with 1x CODEX buffer with ddH20, bottle 2 was filled with DMSO and bottle 3 was filled with ddH20. To start, the antibody-stained coverslip was removed from the storage buffer, transferred to an acrylic stage insert, and mounted with a gasket (Qintay) to safely allow for liquid exchange on top of the coverslips without leaking. The Leica microscope was manually programmed using the LAS X software (Leica Microsystems, version 3.7.1.21655) and set to a 15 μm Z-stack with a Z step size of 1.50 and Z slices of 5. The DAPI channel was used for AutoFocus with a precision level one below 100% accuracy and a capture range of 40 μm. Images were subjected to shading correction by following the instructions on the Linked Shading wizard. Imaged region width and height varied according to the sample size and ranged from 5.4mm to 6.0mm, respectively. Tile overlap was 10%. XY resolution was 324.8 nm/px and Z Pitch 1.499 μm/z. Project settings for imaging collection were: TIFF, uncompressed, and 16-bit.

### Transfer data and image processing

Following image acquisition, raw TIFF images were processed through the standard data transfer protocol using the CODEX Instrument software to organize TIFF files into specific folders based on imaging cycles and regions and rename them according to the format Region_Tile_Z-Plane_Channel.tif. These organized files were subsequently processed using Singer software (Version 1.1.0) from Akoya Biosciences. Singer software performs cycle alignment to compensate for spatial drifts occurring between imaging cycles by referencing the DAPI nuclear stain channel. Additionally, it stitches image tiles together while eliminating overlaps, conducts background subtraction utilizing blank imaging cycles without fluorescent oligonucleotides, and applies Extended Depth of Field (EDoF) processing to merge multiple focal planes into a single, focused image. The software generates a pyramidal qpTIFF image by stacking and compressing all Z-plane, channel, and cycle TIFFs, significantly reducing the file size for subsequent analysis. Processing was carried out using default configuration parameters, with the following functions activated: Extended Depth of Field (EDoF), shading correction, distortion correction, tile normalization, cycle alignment, stitching, and background subtraction. Intermediate TIFF files were saved post-stitching under high-quality performance settings. Additionally, the stitched TIFF images produced by Singer were converted into 16-bit pyramidal OME-TIFF images using the pyvips Python package.

### Cell segmentation

For cell segmentation, we used the QuPath software (version 0.3.2) with deep-learning-based StarDist extension (version 0.3.2)^47^. The StarDist model was retrained using the TissueNet dataset. The segmentation was applied to the DAPI nuclear stain. A probability (detection) threshold of 0.5 was used. Nuclei expansion was implemented with a 5.0-pixel increment, and a cell constraint scale of 1.5 was utilized to approximate the detection of all cells. Intensity and shape features for detected cells were also measured by using the QuPath software. As a result, for all images, tab-separated value (tsv) files were generated. The tsv files were subsequently employed for downstream analysis.

### Hierarchical cell clustering, cluster validation and annotation

Whereas CODEX can support the quantitative analysis of multiple markers in intact tissue samples^18, 24, 48^, its use suffers from technical difficulties, such as signal overlap that can occur between nearby cells, thereby interfering with proper cell-type identification^49^. To overcome these limitations, we developed a hierarchical cell clustering method that leverages ANN classifiers and z-score normalization together with an over-clustering step using K-means. These normalization techniques and unsupervised clustering algorithms have been shown by others to be most effective in mitigating noise-induced clustering artifacts^50^ and are considered as best practices for CODEX analysis. After segmentation, tsv files of each sample were individually imported into QuPath and used to identify single-stained CD140a^+^ tumor cells and cleaved caspase 3^+^ before performing unsupervised cell clustering. To train the classifiers, the train object classifier found on the object classification tab was used to manually select at least 100 different negative and positive CD140a^+^ cells. QuPath’s implementation of the OpenCV multi-layer perceptron was used as a classifier while whole-cell protein intensity values were used as features. These same steps were also used to train the classifiers for cleaved caspase 3^+^ cells. After this, tsv-files were exported and subjected to R and Python scripts, where all cells that were negative for CD140a and cleaved caspase 3 from both control and α-CTLA-4/α-PD1 samples were pooled together, and each marker intensity was subjected to Z-score normalization. As shown in Hickey et al^50^, this normalization method reduces clustering artifacts associated with low signal or high background. Importantly, to circumvent batch effects and run-to-run variations, we normalized across pooled runs only after marker-level QC and z-score normalization, and we verified consistent staining patterns across days and conditions. After this, the normalized dataset was subjected to two rounds of K-means clustering based on the expression of lineage-defining markers but excluding phenotypic markers (Ki67, GZM-B, ICOS, PD-1, Lag-3, TIM-3, PD-L1). This strategy was chosen because of the published recommendations, indicating that phenotype annotations should be avoided at the clustering step and that over-clustering enhances the separation of cell types often confounded by noise into distinct clusters. The Lloyd algorithm was used for k-means clustering with k=60 for both rounds of clustering, of which the known immune cell types were identified, annotated and merged using average cluster profiles. The second round of clustering was performed only with the clusters that we couldn’t properly annotate in the first round, given the low expression or mixed signal intensities of mutually exclusive lineage-defining markers. This produced another 44 clusters, of which the known cell types were combined with the first round of clustering, resulting in the final annotation of 13 known cell types, plus a population of unidentified immune cells and a population negative for all immune lineage-defining markers labeled as other. These 13 cell types were then visually verified by overlaying the cells of each cluster onto the stitched images in QuPath.

### T-distributed stochastic neighbor embedding (tSNE) dimension reduction

To visually confirm the distinct identity of the immune cell populations, 100,000 immune cells were sampled across both groups and subjected to tSNE dimension reduction^51^, where the high-dimensional expression profiles of the cells were transformed into a two-dimensional space^51^.

### Cellular neighborhood (CN) analysis

To identify CNs in T3 tumors, we applied the “Sliding Window Mapping Method” developed by the Nolan Lab^20^, which uses a custom k-nearest neighbor approach available via GitHub (github.com/nolanlab/NeighborhoodCoordination). We set the window size to 20 neighbors, consistent with parameters from the original study. Each window was represented as a vector containing the frequency of 13 defined cell types. Following window generation, CNs were identified using the MiniBatchKMeans clustering algorithm (scikit-learn v1.2.1), with the number of clusters (K) set to 20 and neighborhood size to 7. This configuration was chosen based on prior CODEX studies to balance resolution and biological interpretability. Higher values of K tended to split known CNs without revealing new biological patterns, while smaller values merged distinct tumor, boundary, and immune compartments. Similarly, increasing the neighborhood size beyond 7 did not yield novel CNs but instead divided cell types into redundant or overly similar groups. We then calculated the log2 fold change (log2 FC) of cell-type enrichment within each neighborhood relative to the tissue background. The log2-transformed fold change values were then visualized as a heatmap using the sns.clustermap function from Python’s Seaborn library^52^. We report full parameterization and code to allow replication.

### Immunity-promoting sub-neighborhood (IP-SN) analysis and spatial correlation

We also applied the “Sliding Window Mapping Method” specifically to the cells encompassing the immunity neighborhood. For this analysis, CD4^+^ and CD8^+^ T cells were further annotated based on their z-scaled expression values for GZMB and Ki67. This refined data was later used for sub-neighborhood analysis within the IP-CN6. The resulting heatmaps for each time point were used for correlation analysis. The expression column profiles of GZMB^+^ T cells and Ki67^+^ T-cells were utilized to measure the Pearson correlation between them.

To quantify co-localization of GZMB⁺ and KI67⁺ T cells within the IP-CN6, we examined the IP-SNs heatmap for each time point and annotated IP-SNs that were simultaneously enriched for both populations. For each cell type, we then calculated the percentage residing in these co-enriched IP-SNs. Analogously, we identified IP-SNs co-enriched for cDC1 with GZMB⁺ T cells and for cDC-1 with KI67⁺ T cells. For cDC1, the percentage residing in each set of co-enriched IP-SNs was computed relative to the total cDC1 count across all IP-SNs using the same approach.

### GZMB^+^ and Ki67^+^ distance calculations

We performed a spatial distance analysis between GZMB⁺ CD8⁺ T cells and Ki67⁺ CD8⁺ T cells relative to tumor bulk cells to determine the average shortest distance between T cells and tumor cells. We present full distance distributions **(Figure 7D)** and summarize fractions above/below 350 μm as a mid-distribution reference chosen at d10 when Ki67⁺ CD8⁺ T cell split approximately 50:50 **(Figure 7E)**. Conclusions do not rely on a single cutoff, as the distributions separate over time. This analysis was conducted using the spatial_distance function from the scimap Python package.

### Triads frequency analysis and T cell phenotype within triads

To quantify the frequency of cDC1 or IFNγ–activated macrophages participating in two-cell (dyads) and three-cell (triads) clusters involving APCs, CD4⁺ T cells, and CD8⁺ T cells, we defined a fixed 10μm radius centered on the centroid of each APC’s segmentation mask. This radius served as a proxy for potential synaptic engagement, in line with previous spatial analyses. However, it is important to emphasize that this approach does not confirm direct membrane–membrane contact. Additionally, given the use of 8μm tissue sections and EDoF technology, spatial projections may include vertically offset cells, potentially leading to overestimation of physical interactions within the 10 μm threshold. To specifically identify triadic clusters, we first identified all neighbors within the fixed radius using the frNN function from the dbscan R package. We then classified these clusters as dyads if they contained two cells (one query APC and one T cell) or as triads if they comprised among neighbors CD4 and CD8 T cells with exclusion of regulatory T cells. Alternatively, we further stratified these triads by T cell phenotype within each triad. By extracting high-resolution phenotype data for all neighbors within defined triads, we quantified the proportions of T cells that were GZMB^+^, Ki67^+^, or double negative (GZMB⁻Ki67⁻) over time.

### Proximity density score

To quantify the frequency of interactions between specific cell types, we calculated Proximity Density. This metric assesses the likelihood of interaction between two cell types relative to their abundance within the tissue. Specifically, Proximity Density reflects the ratio of observed interactions between two cell types to the expected number of interactions based on their respective population sizes. We utilized the scimap Python package available at ^53^ to compute Proximity Density scores for the following cell type pairs: cDC1 cells and Ki67^+^ T cells.

### Permutation test

To identify significant co-localizing or avoidance behaviors between different cell types, we utilized the spatial interaction function from the scimap Python package. We conducted permutation tests of single-cell interactions following the methodology previously described^29, 30^. In our analysis, cells located within a 6-pixel radius (6 μm) were considered to be co-localizing. Significant co-localizing or avoidance behaviors were defined by a P-value threshold of less than 0.01. Briefly, we defined a co-localization zone with a 6 μm radius extending from the nucleus of any one cell to the outermost edge of an adjacent cell. Cells within this radius were considered to be co-localizing. Then, to identify non-random patterns of co-localization, we performed 1,000 permutations of the cell labels while fixing their x and y coordinates, allowing us to differentiate between co-localization and avoidance behaviors. Significant co-localization among different cell types was defined as having a p-value of less than 0.01

### Longitudinal proximity between CNs

To analyze the collective longitudinal actions of the cell types comprising each CN, we labeled cells according to their respective CNs and performed a 20-nearest-neighbor analysis on each cell across multiple time points. This approach quantified how the spatial proximity between CNs changed longitudinally. To assess the significance of the inter-CN interactions, we used a t-test followed by Benjamini-Hochberg adjustment for multiple hypothesis testing for each CN-to-CN interaction pair.

### Network of communication between CNs

To characterize the tissue assembly rules governing CN communications, we first identified the cells that were located at the interfaces between adjacent CNs. To do this, we identified 20 nearest neighbors for each cell, and cells with at least 30% of their neighbors belonging to an adjacent CN were defined as “interface cells”. We then performed the permutation test by using the spatial_interaction function from the scimap Python package to assess the significance and level of interaction between different CN interfaces. This allowed us to build a network of interactions between the different CN-interfaces. The results were represented as a network plot, where the node sizes represent CN frequency in the tissue at a particular time point, and connection arrows between nodes indicate the strength of interaction between interfaces, ranging from weak (light gray) to strong (black).

### Sorting of scRNA-seq samples

For sorting of tumor cells and intratumoral immune populations from d10 T3 tumors, single-cell suspensions were generated from enzymatically digested tumors. CD45⁺ immune cells were first enriched using CD45 magnetic microbeads (Miltenyi Biotec, cat. no. 130-052-301). Enriched CD45⁺ cells were incubated with Fc block (anti-CD16/32; 500 ng) for 5 minutes at room temperature, followed by staining with an antibody cocktail containing anti-CD45, anti-CD4, anti-CD8, anti-XCR1, anti-CD11c, and anti–MHC class II antibodies (BioLegend; 1:200 dilution) in 100 μL staining buffer (PBS supplemented with 2% FCS) for 30 minutes at 4 °C. In parallel, the CD45⁻ fraction from the same single-cell suspension was stained with anti-CD140 antibody to isolate T3 tumor cells. Cell sorting was performed on a BD FACSAria II (BD Biosciences). The purity of all sorted populations used for single-cell RNA sequencing exceeded 96%, as assessed by post-sort analysis.

All sequencing samples were then exported and combined for further single-cell analyses in Seurat^54^ and scanpy^55^. To visualize cells from multiple collection batches, we selected for the top 8,000 highly variable genes (HVGs) using the ‘sc.pp.highly_variable_genes’ method with ‘seurat_v3’ option and correcting for collection batch, and we performed PCA on logarithmized, normalized expression and UMAP dimension reduction using the top 30 principal components and neighborhood size of 10. The integrated object was also subset to tumor and immune cells, based on collection batch annotations (i.e., CD45 status prior to sequencing), to create the dataset for cell type annotation and additional downstream single-cell analyses.

### Annotation of scRNA-seq cells

Major immune cell types were first identified by manual annotation of differential gene expression between clusters in the immune cell subset based on known markers and signatures^14^. We initially performed clustering using the Phenograph method implemented in scanpy^56^ with k=300 and resolution=1.0 and annotated cell type groups based on major lineages (macrophage, CD8^+^ and CD4^+^ T cell, Treg, natural killer, dendritic cell, neutrophil) and cell cycle status. We then subset the object to macrophages and re-performed Phenograph clustering with k=200 and resolution=1.0 and annotated macrophage subtypes based on known markers and signatures^14^. After functional annotating macrophages and T cells, we selected cells with high-confidence and relevant immune subtyping for further analysis.

### Differential gene expression and gene set enrichment methods

To identify differentially expressed genes (DEGs), we performed standard T-tests on logarithmized, normalized expression for each gene marker and Benjamini-Hochberg P-value correction, implemented in scanpy as the ‘sc.tl.rank_genes_groups’ method. Vlad to fill out gene set enrichment details.

### ARCADIA overview

ARCADIA is a dual variational autoencoder (VAE) framework that integrates scRNA-seq data with spatial proteomics data^32^. ARCADIA first identifies and matches archetypes between modalities, defined as common extreme phenotypic states, rather than relying on feature-level linkage for integration. It then specifically establishes “anchors”, which are high-confidence cell state archetypes that guide the alignment of all cells between modalities.

First, we denote *N* cells and *G* genes in scRNA-seq data (*X^RNA^*) and *M* cells and *P* protein features for CODEX data (*X^Protein^*). We describe ***z^RNA^*** and ***z^Protein^*** as low-dimensional latent representations for each data modality, and ARCADIA entangles these latent spaces using a multi-objective function that ensures gross- and fine-grained details are matched properly across modalities. The generative process for gene expression counts X ^RNA^ is modeled by a zero-inflated negative binomial distribution, ZINB(μ_ng_, θ_g_, π_ng_), where mean expression μ, gene dispersion θ, and dropout probability π are parameterized by neural networks. Simultaneously, protein expression X_mp_^Protein^ is modeled by a Gaussian distribution *N*(μ_mp_, σ²_p_), where mean expression μ and variance σ² are parameterized by separate neural networks.

To learn entangled latent representations, the framework optimizes a joint objective function that combines reconstruction loss (Evidence Lower Bound or ELBO) with geometric alignment constraints, incorporating known biological states and extracted anchors. The total loss function is defined as L_total_ = Σ (λ_0_ * L_ELBO_ + λ_1_ * L_struct_) + λ_2_ * L_anchors_ + λ_3_ * L_cross-modal_, evaluated for both scRNA-seq and CODEX and where hyperparameters λ_0_, λ_1_, λ_2_, and λ_3_ are positive weights controlling the relative contribution of each objective component as implemented in the original study^32^. We minimize this total loss using stochastic gradient descent to optimize for the encoder and decoder parameters. Additional mathematical details are available in ^32^.

The objective function consists of four key components. First, Reconstruction Fidelity (L_ELBO) uses the Evidence Lower Bound to ensure accurate reconstruction of the original features for each modality. Second, Structure Preservation (L_struct) maintains the internal structure of major cell types within each modality, ensuring that distinct cell types and major patterns remain separated. Third, Cross-Modal Alignment (L_cross-modal) uses Maximum Mean Discrepancy (MMD) to align the global distributions and major patterns of cell types across the two modalities. Finally, Anchor Alignment (L_anchors) penalizes latent distances between biologically matched archetype anchors. This term is responsible for retaining subtle intra-cell type patterns, particularly those driven by the spatial environment that cause phenotypic differentiation within cells of the same type.

### scRNA-seq and CODEX data preprocessing for ARCADIA integration

We applied ARCADIA to integrate scRNA-seq and CODEX data from day 10 T3 tumors harvested from α-CTLA-4/α-PD-1 and control mice, and we focused our analysis on immune cell states. To avoid biases from the relative number of cells recovered in each assay, we integrated the entire scRNA-seq dataset with one representative day 10 control and one representative d10 α-CTLA-4/α-PD-1-treated CODEX sample. The RNA count and protein expression values for each dataset were normalized and pre-processed for ARCADIA input as described previously^32^. Specifically, we selected 2,000 HVGs and logarithmized and normalized scRNA-seq prior to archetype matching (scRNA-seq remained as count data for dual VAE training); for CODEX data, we retained all protein markers and normalized data using median absolute deviation and z-scoring. In addition to cell-intrinsic protein expression, we incorporate spatial protein expression features for each CODEX cell by averaging neighborhood expression from the 20-nearest neighborhood graph (the default setting). We used CN assignments computed as described above for the α-CTLA-4/α-PD-1-treated and control CODEX sample. Cell annotations were harmonized between scRNA-seq and CODEX datasets for d10 samples, and only immune cell subtypes present in both datasets were aligned.

### Archetype alignment and integration by dual VAE training

We defined archetypes for each data modality using Principal Convex Hull Analysis (PCHA) on the preprocessed feature spaces. To account for phenotypic extremes due to α-CTLA-4/α-PD-1 treatment effects, we computed *k* archetypes for each α-CTLA-4/α-PD-1-treated or control sample (optimal *k* chosen between 7 and 13 based on cross-modal matching quality), merged archetypes across samples within each modality, and then aligned archetypes across modalities using the Hungarian algorithm on cell-type composition profiles. Anchors to guide ARCADIA integration were defined as high-confidence cells with dominant archetype weights in the top 5% of each archetype’s distribution.

We trained the dual VAE architecture for 500 epochs using the AdamW optimizer with a learning rate of 1e-3 and a batch size of 1,024. Both VAEs were instantiated with latent dimensionality of 60 and 3 hidden layers for the encoder and decoder neural networks; the RNA VAE used 1,024 nodes per hidden layer while the protein VAE used 512 to account for lower dimensionality. The model optimized the previously described objective function incorporating modality-specific reconstruction, within-modality cell type clustering, anchor matching, and cross-modal cell type alignment losses. By default, the relative contribution of each objective component (each *λ_i_* scalar) is automatically tuned so each loss term scales proportionally. Other hyperparameter values were chosen based on clustering and integration metric performance^32^. In optimizing the joint objective function, each modality’s latent space becomes entangled (**Figure S7C**) which aligns immune cell states from both datasets into a unified coordinate system – the ARCADIA entangled latent space – which could then be used to predict spatial niche demographics for scRNA-seq cells (**Figure S7D).** When visualizing cells from both modalities in ARCADIA space, CODEX cells are subsampled to 30% to better match scRNA-seq cell counts.

### Analysis of spatially dependent differential gene expression

Using the entangled latent space, which reflects phenotypic similarities and variation based on shared archetypes, we predicted CN labels for each scRNA-seq cell informed by its 10 nearest-neighboring CODEX cells. Subsequently, we performed differentially expressed gene (DEG) analysis between α-CTLA-4/α-PD-1-treated and control cells, for each cell type within different CNs. To focus our analysis, we only examined DEG patterns from cell types assigned to CN6 with adequate representation, for example, CD8^+^ and CD4^+^ T cells in CN1, CN2, CN4 and CN6. DEGs were identified using one-versus-all T-tests on logarithmized, normalized expression and Benjamini-Hochberg P-value correction, with predicted CN labels as the testing groups. The resulting DEGs were individually interpreted (analysis of top 100 statistically significant up- or down-regulated genes based on log-fold-change) and analyzed via gene set enrichment (GSEA). Specifically, GSEA was performed using the *fgsea* R package^57^ on ICT vs control DEGs for cells within CN6 cells or using cells from all CNs. In the main and supplemental figures, CD8 and CD4 T cell marker gene expression patterns across predicted CNs are depicted in using dotplots, where dot size is the mean expression of α-CTLA-4/α-PD-1-treated cells within a CN and sizes are scaled for each marker across CN groups.

### Treatment-dependent cellular interaction inference from scRNA-seq using ContactTracing

ContactTracing identifies conditionally-dependent cell-cell interactions and captures cell type-specific responses to ligand-receptor-mediated interactions^33^. We applied ContactTracing on scRNA-seq from immune cells from d10 α-CTLA-4/α-PD-1 and control samples, using α-CTLA-4/α-PD-1 treatment as the experimental condition parameter. For cell annotations, we used major cell type labels for all immune cell populations except macrophages, for which we used functional cell type labels based on hierarchical clustering and annotation. Other than feature selection, which was not performed to keep the analysis unbiased and transcriptome-wide, we used default parameters and cutoffs as originally implemented. For cell-cell interaction Circos plot visualizations, cell type-specific ligands and receptors were ordered by diffusion components computer on differential expression as in the default implementation. To ensure high confidence in inferred interactions, we only studied interactions with differential ligand expression FDR<0.01, ligand expression in >5% of each donor cell type, ligand effects involving more than one differentially expressed gene, thus highlighting significant interactions with meaningful downstream transcriptional effects across immune cell types. To further focus our investigations, we visualized networks of T cell-APC triad-specific interactions invigorated or abrogated by α-CTLA-4/α-PD-1 (positive or negative ligand log-fold-change).

### Statistical analyses

A *P* value of < 0.05 was considered statistically significant. All statistical testing methods are described in figure legends.

