## Supplementary figures and images for "Immune Checkpoint Therapy Drives Maturation of a Cellular Neighborhood Nucleated by T Cell-APC Triads Enabling Spatially Compartmentalized Tumor Immunity"

### Supplemental figure 1

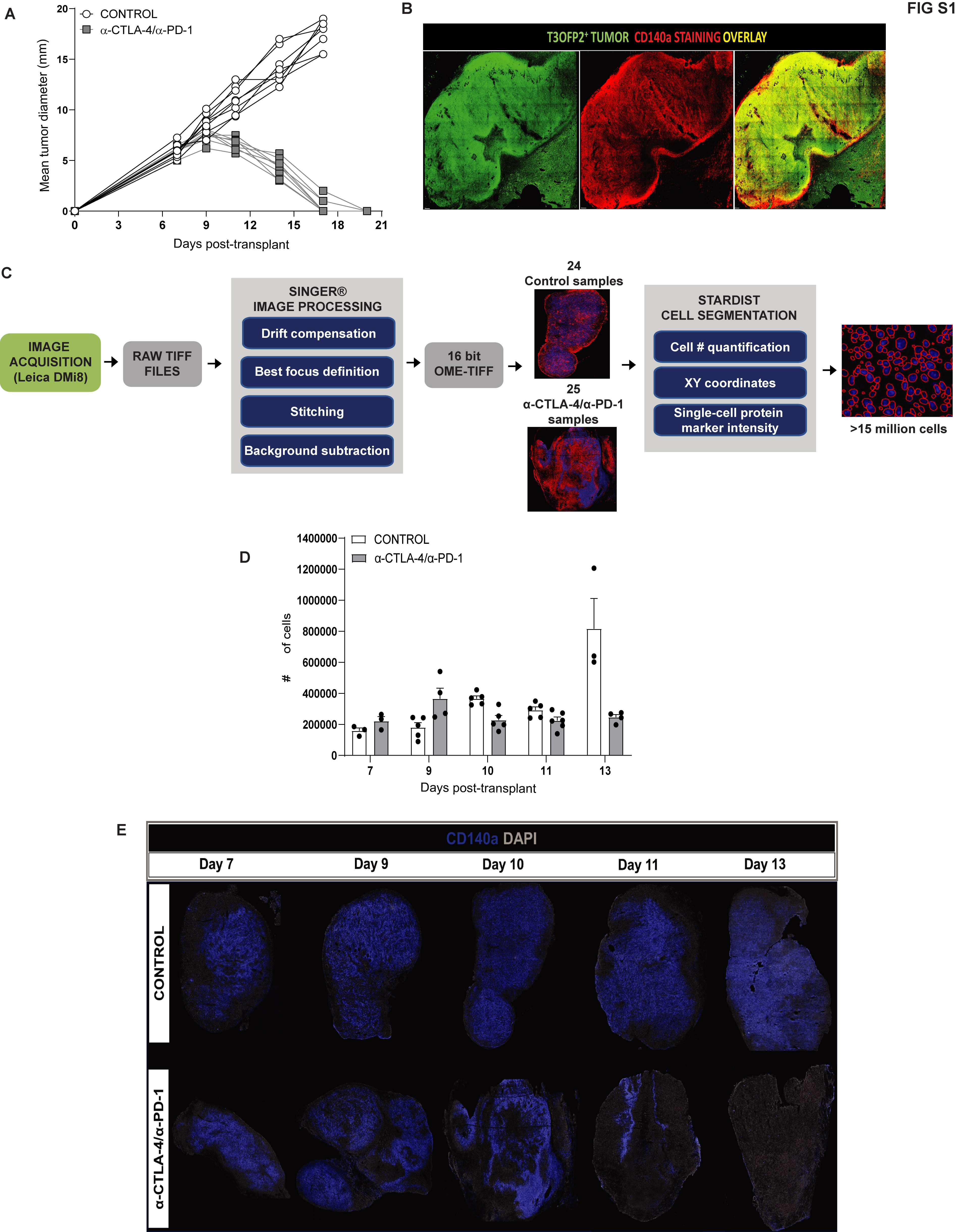

### Supplemental figure 2

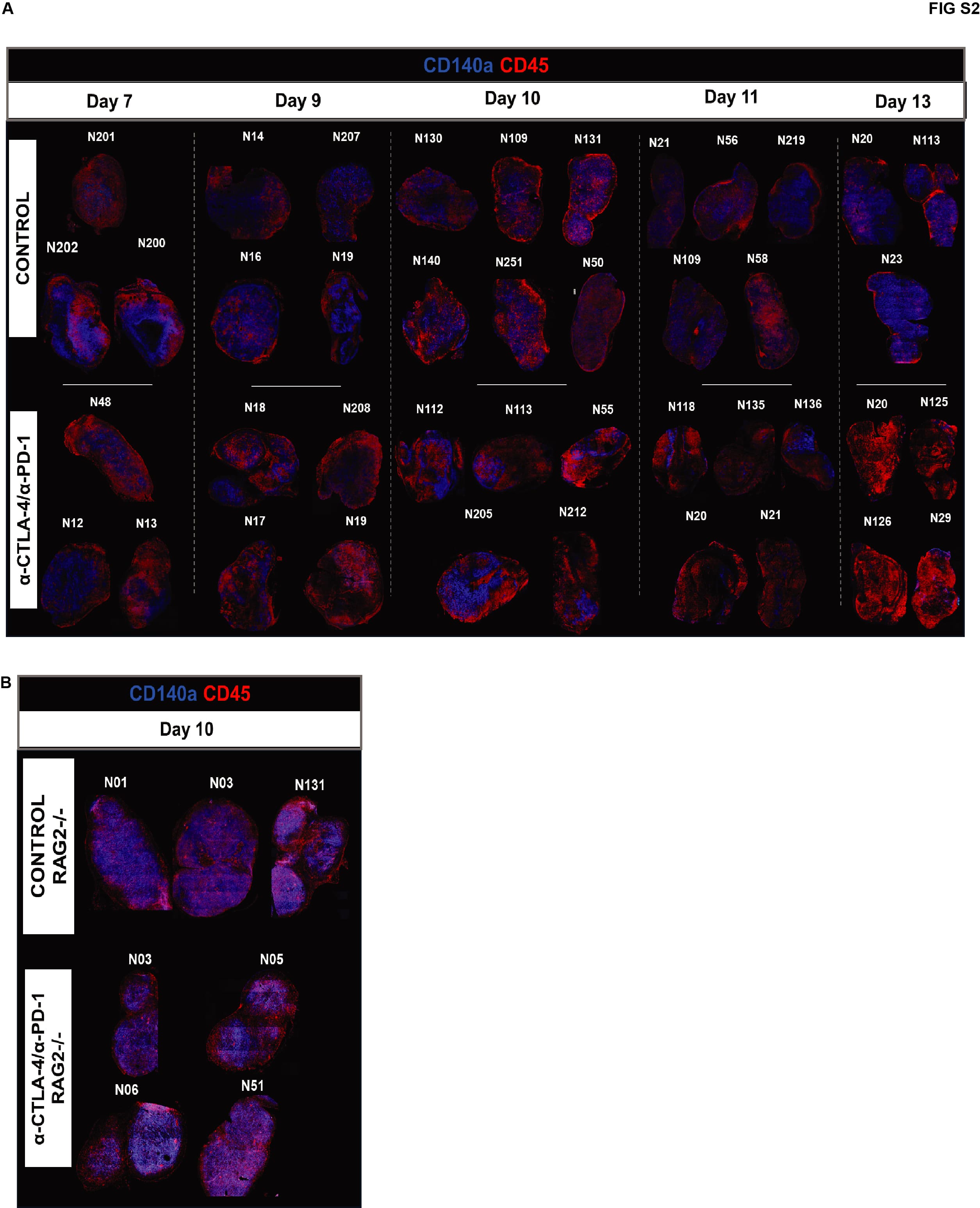

### Supplemental figure 3A

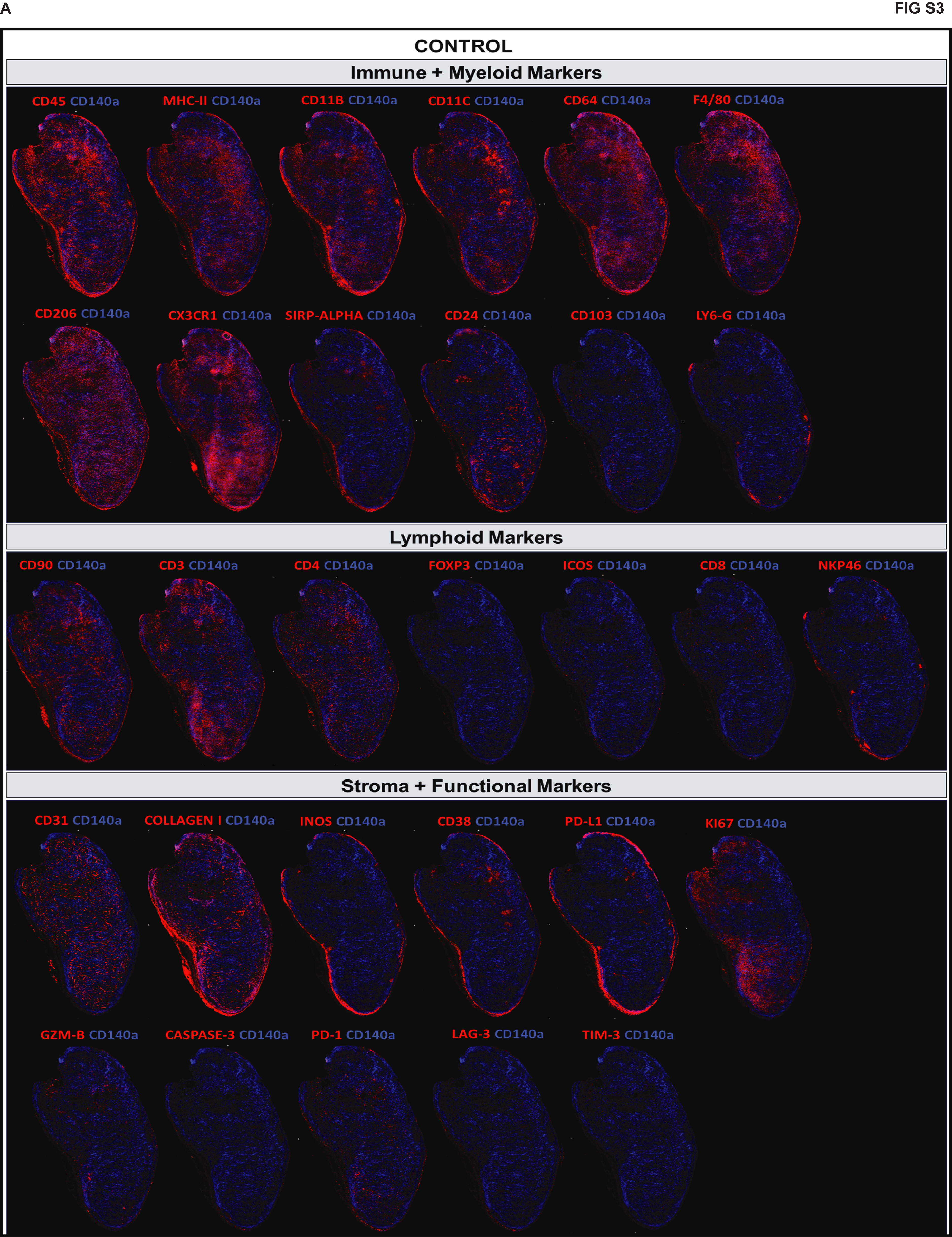

### Supplemental figure 3B

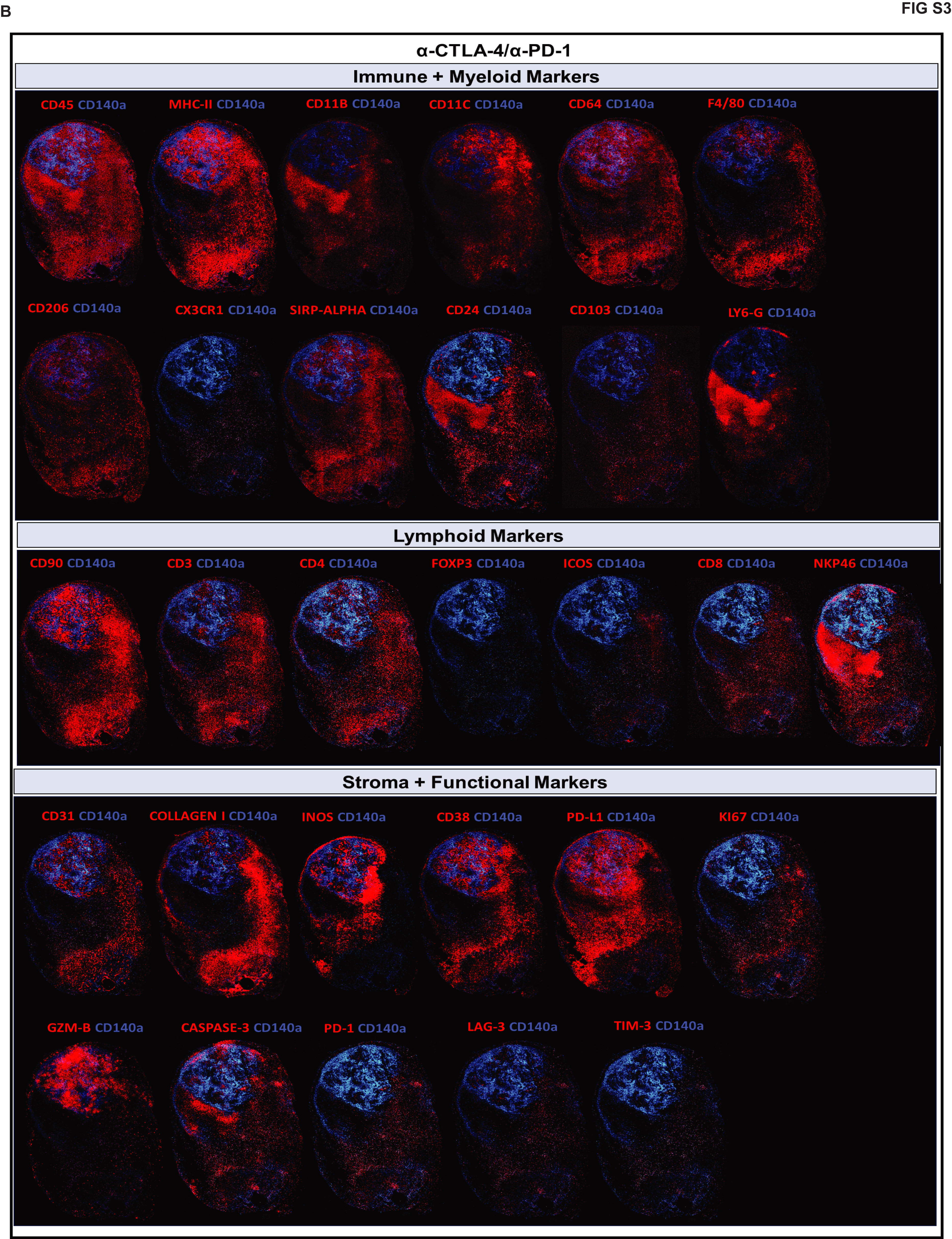

### Supplemental figure 4

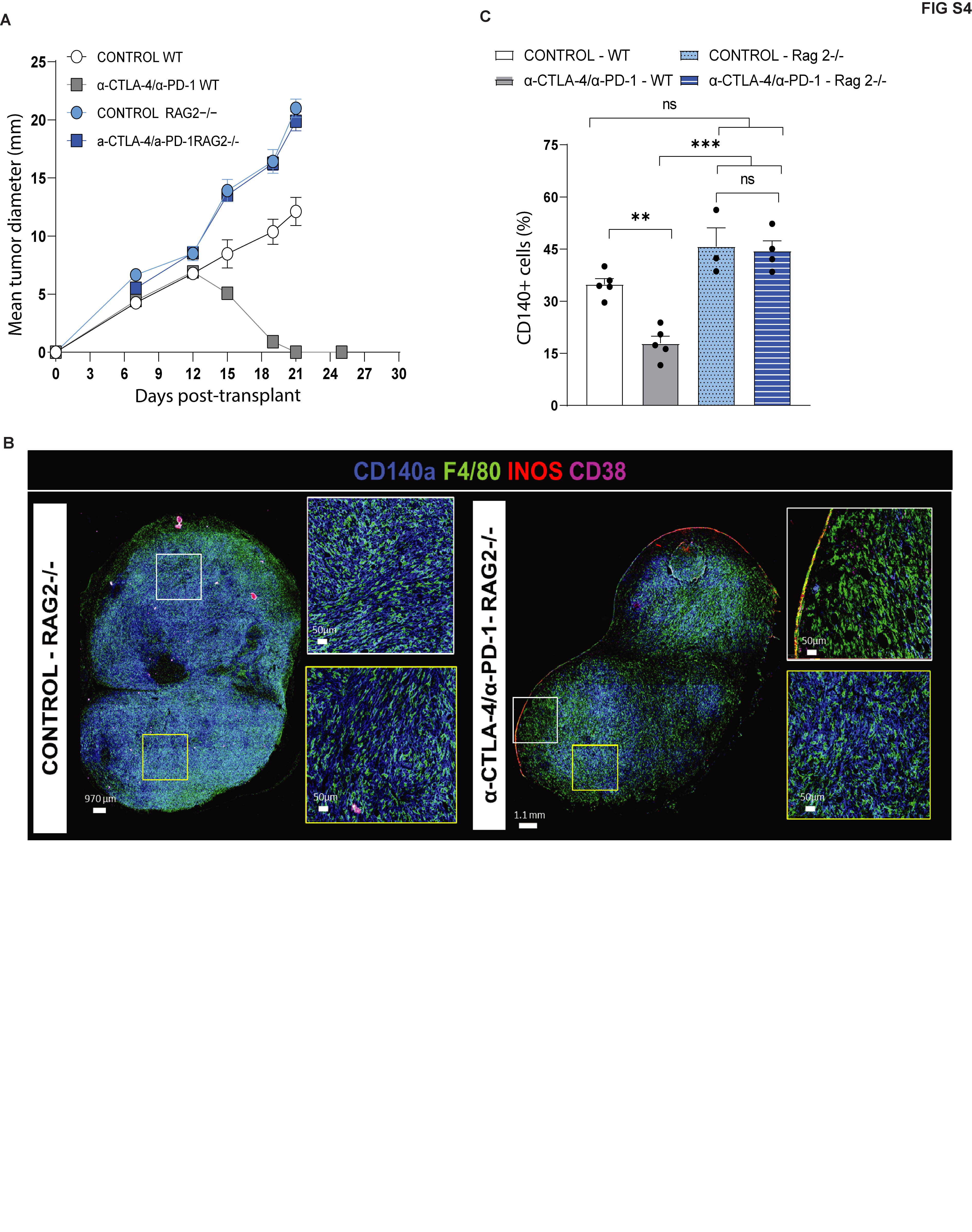

### Supplemental figure 5

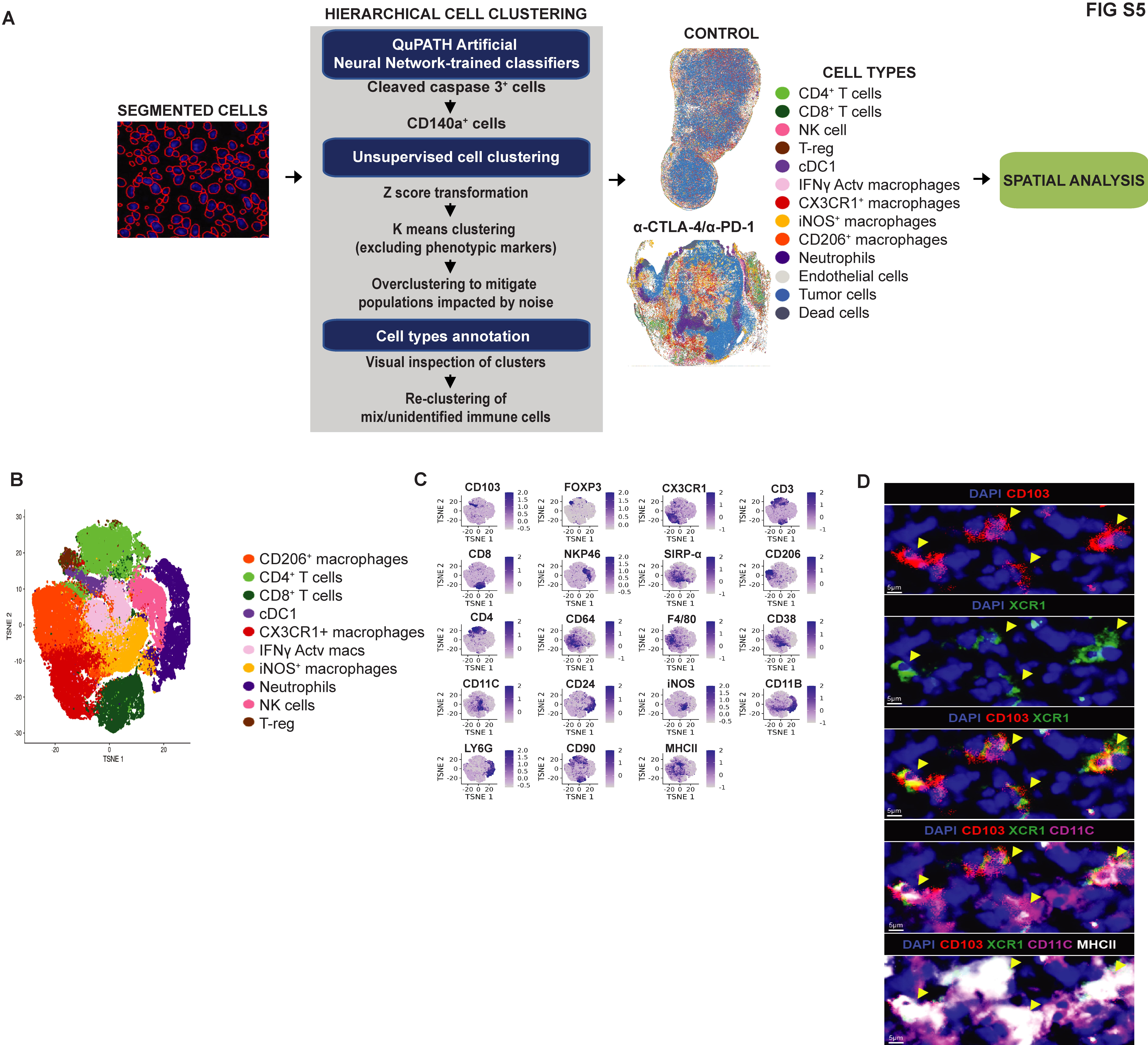

### Supplemental figure 6

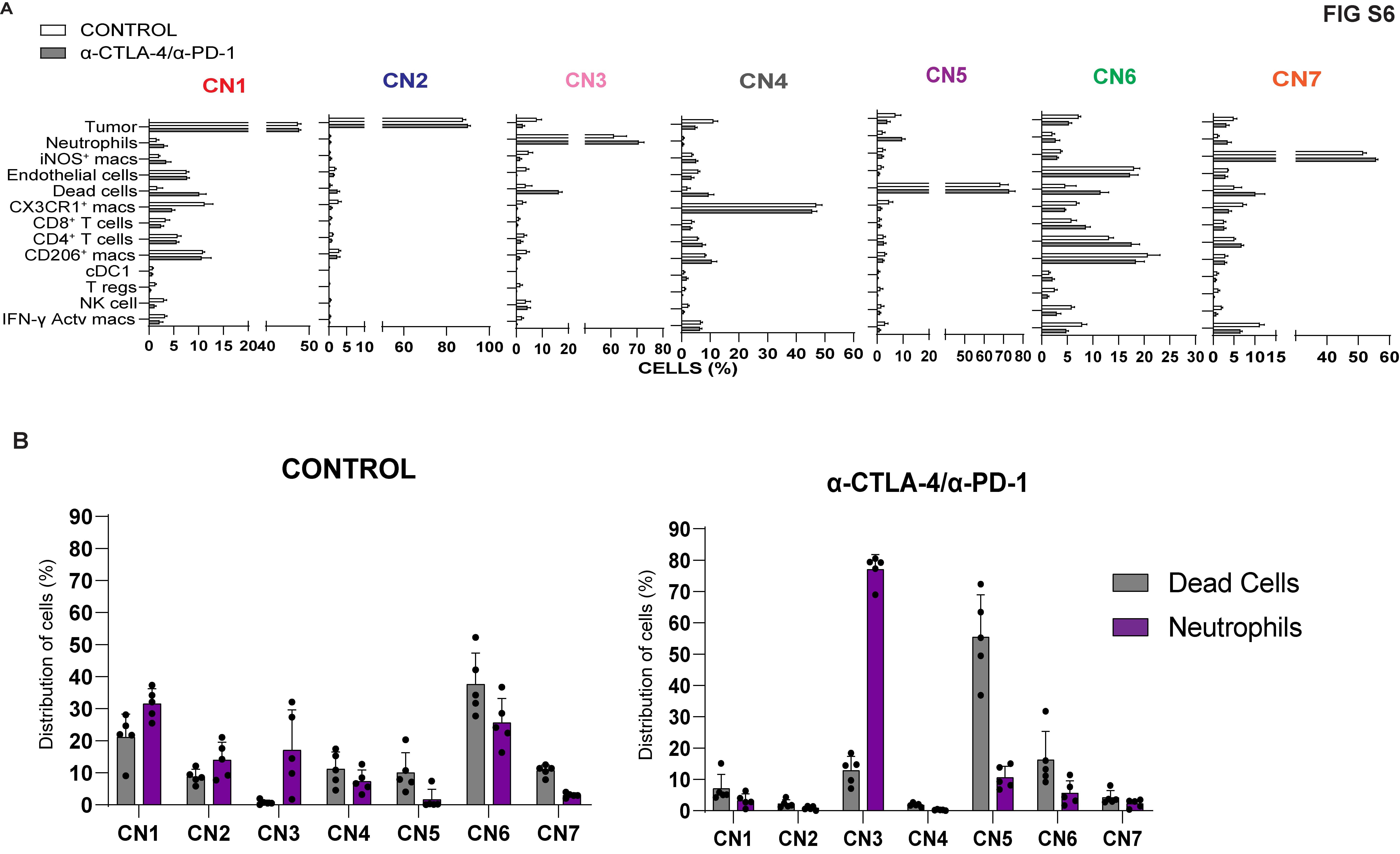

### Supplemental figure 7

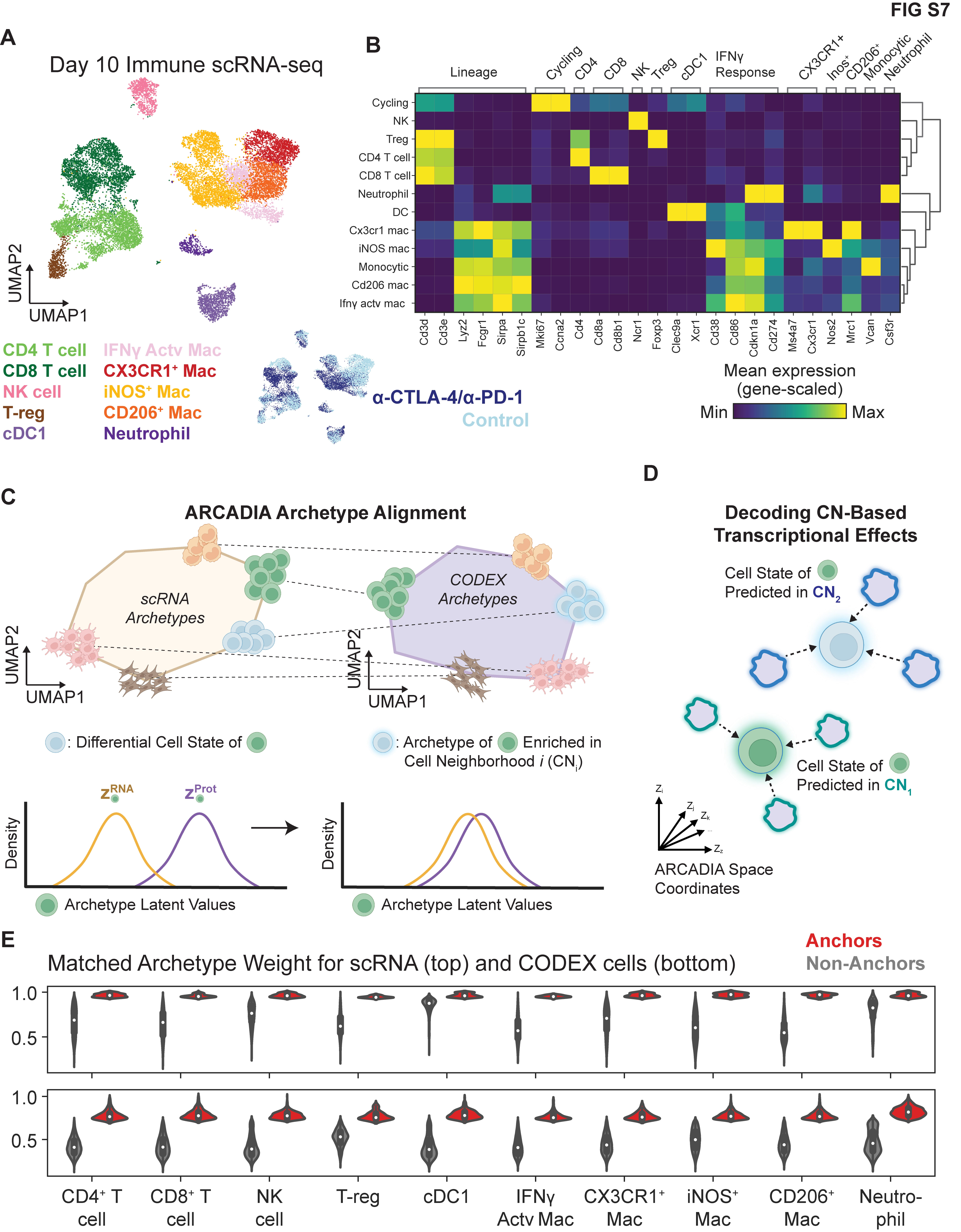

### Supplemental figure 8

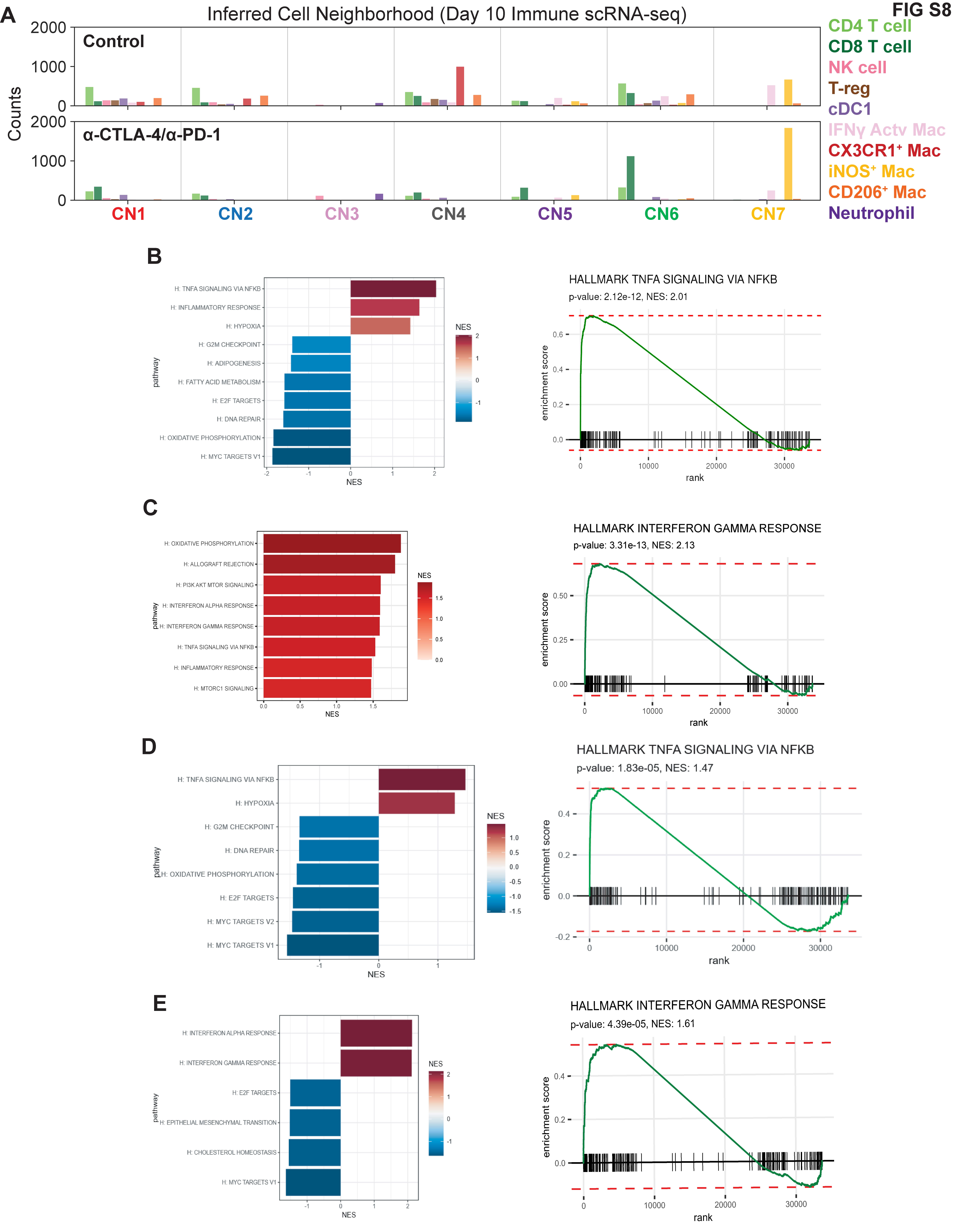

### Supplemental figure 9

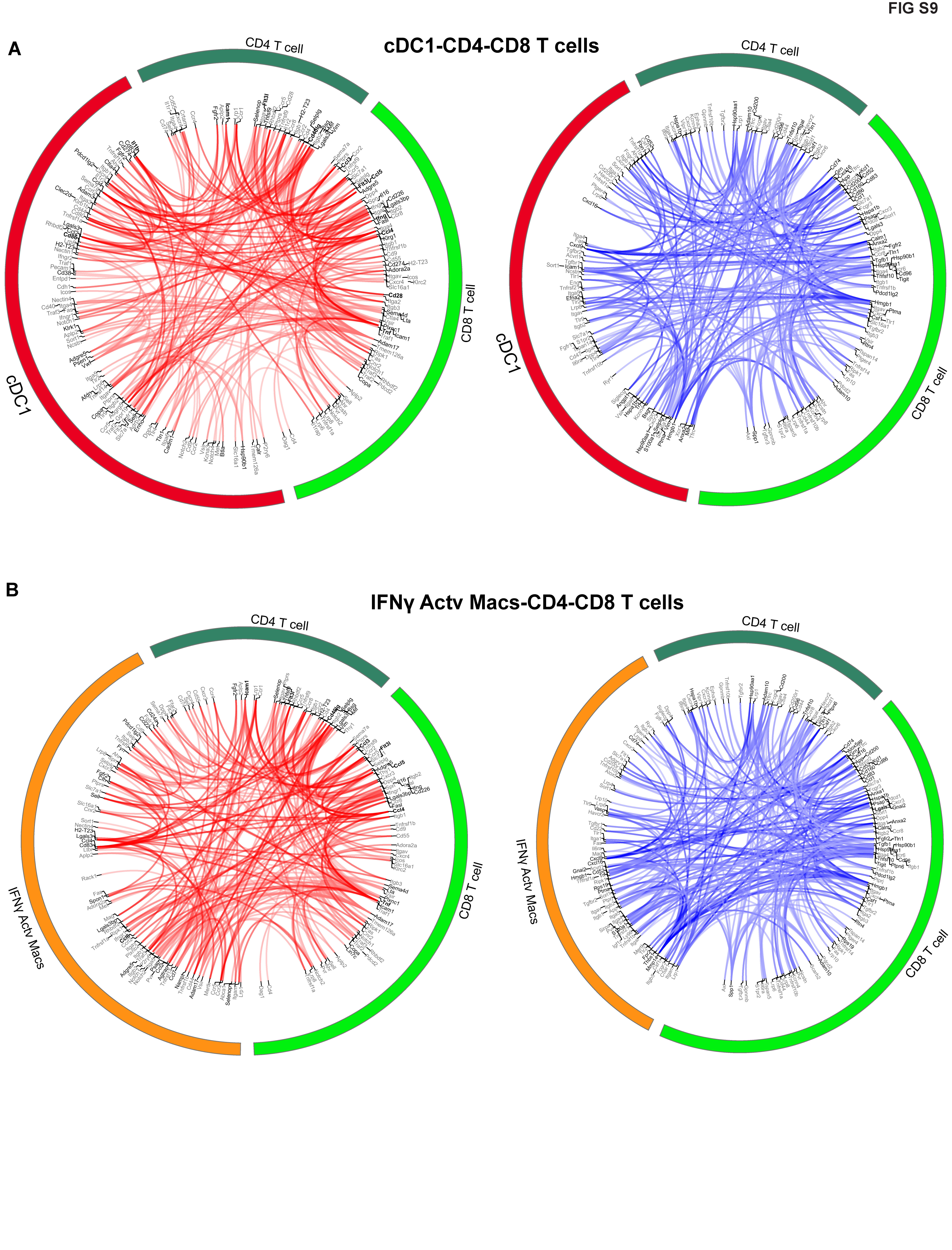

### Supplemental figure 10

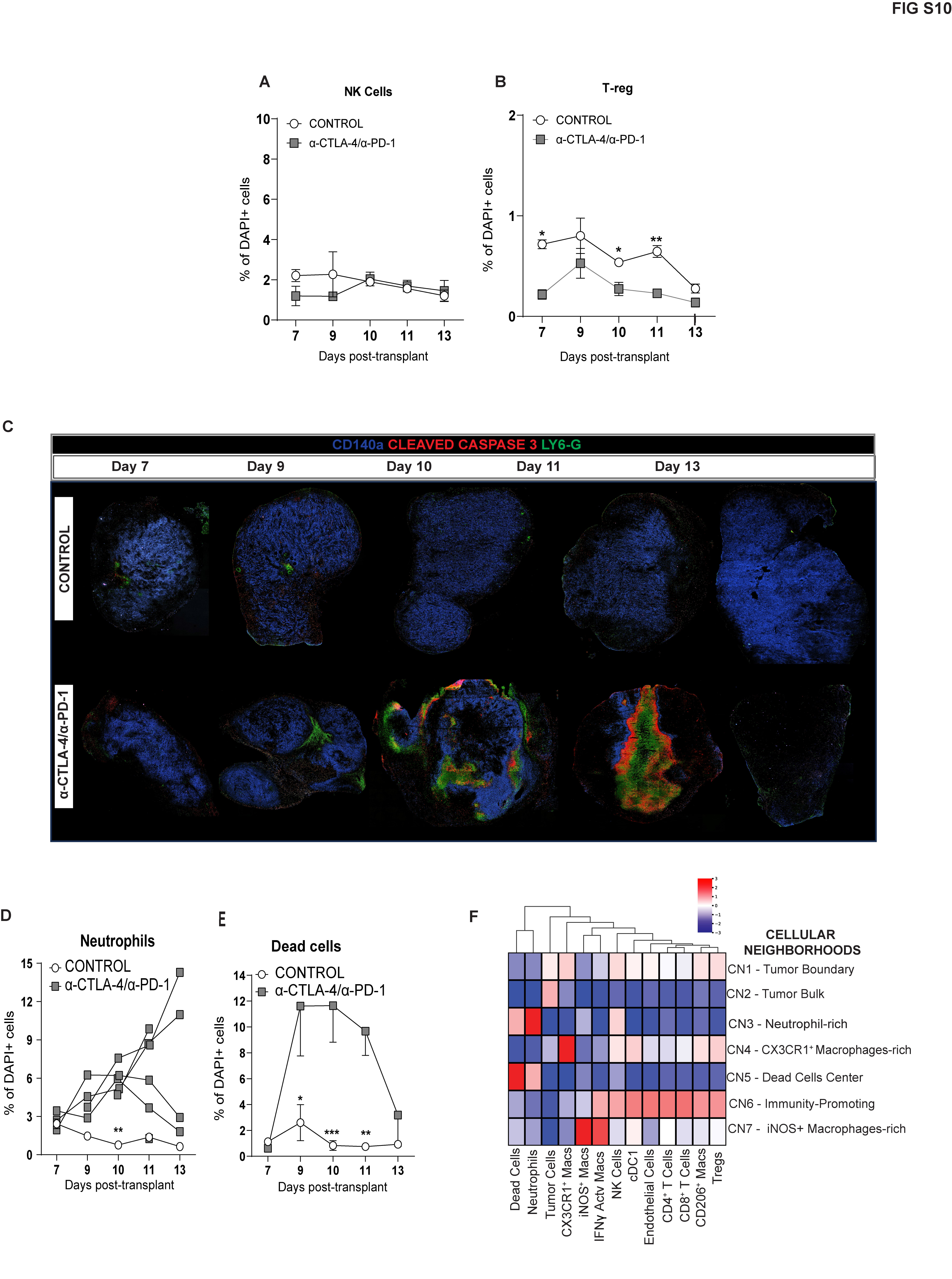

### Supplemental figure 11

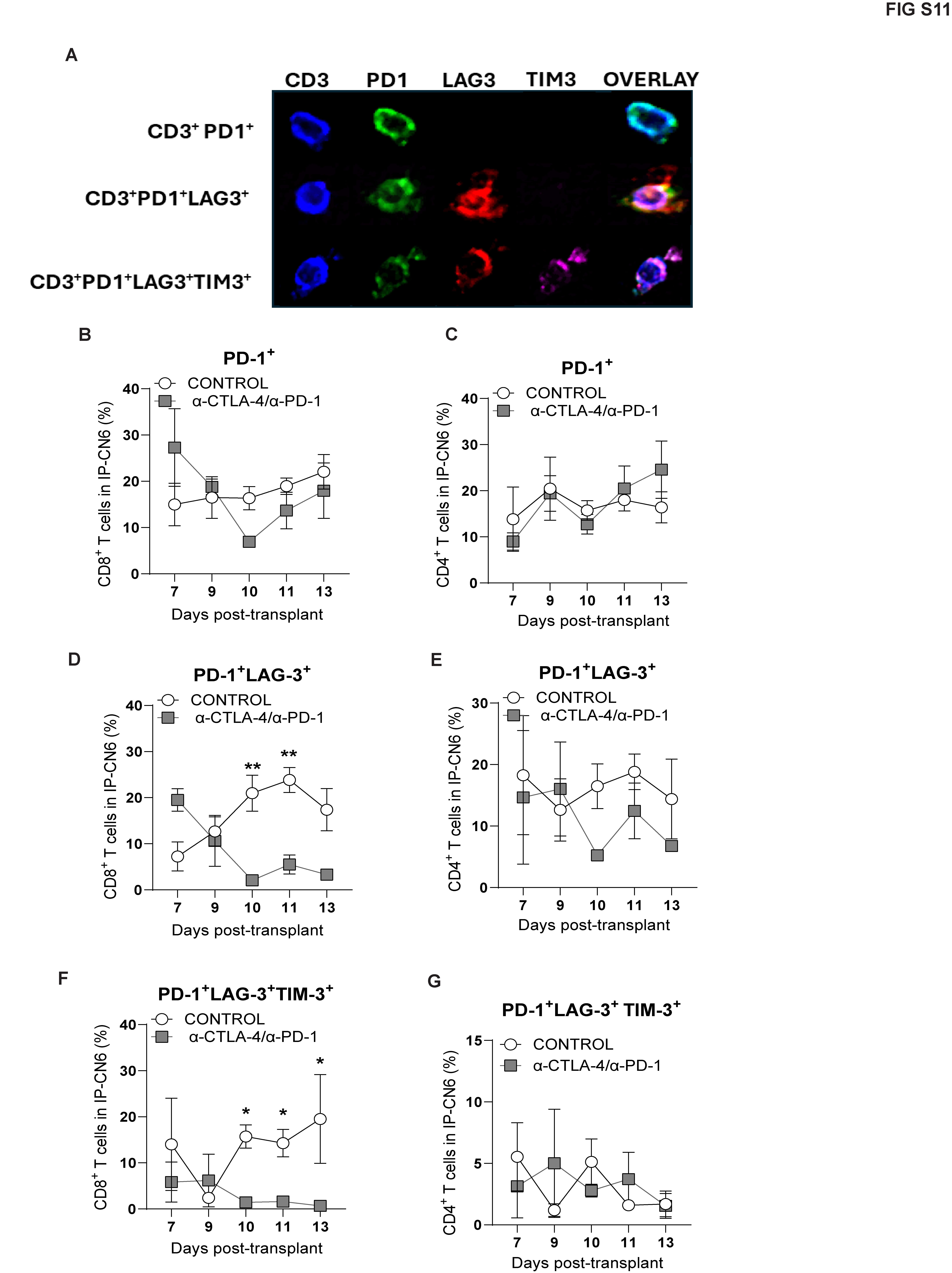

### Supplemental figure 12

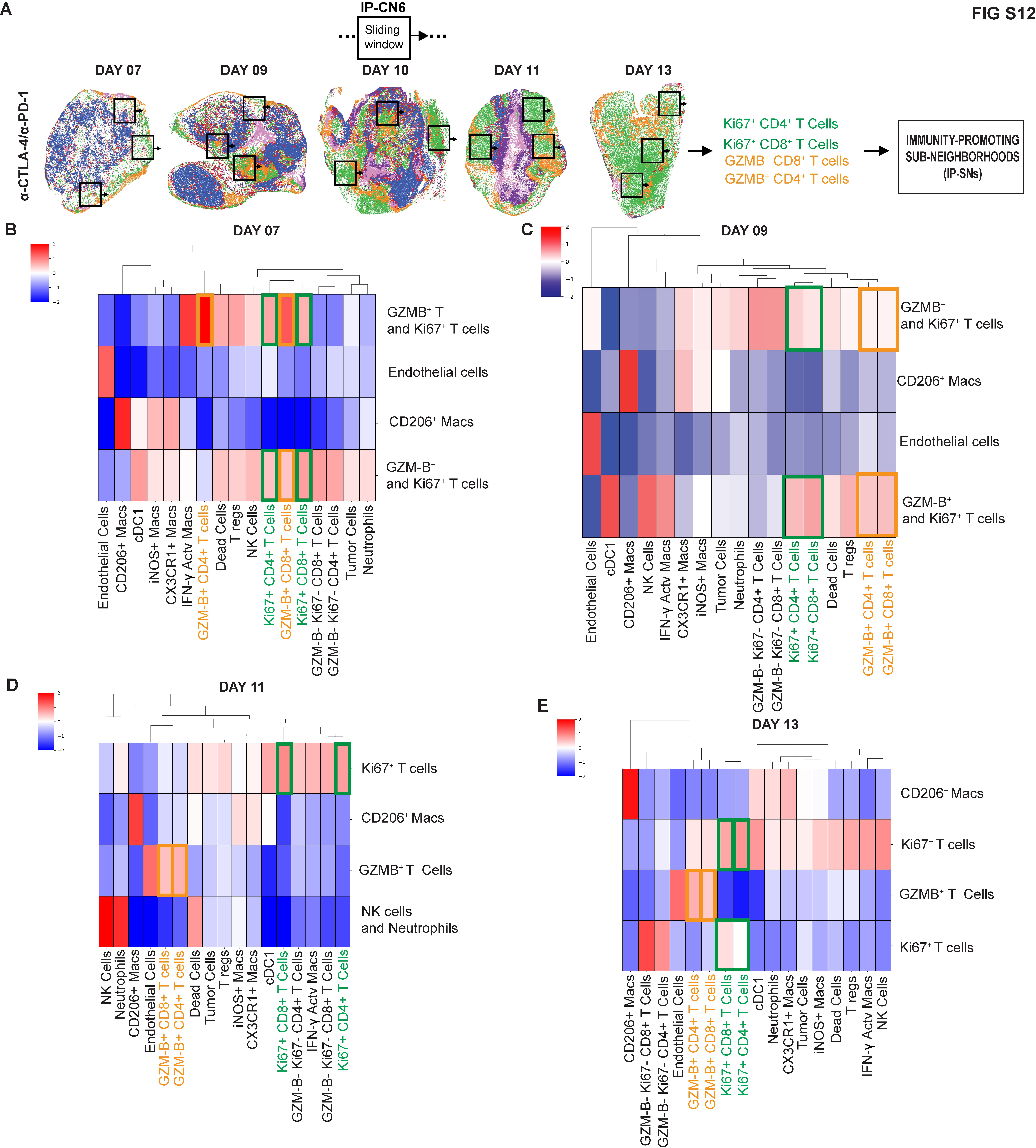

### Supplemental figure 13

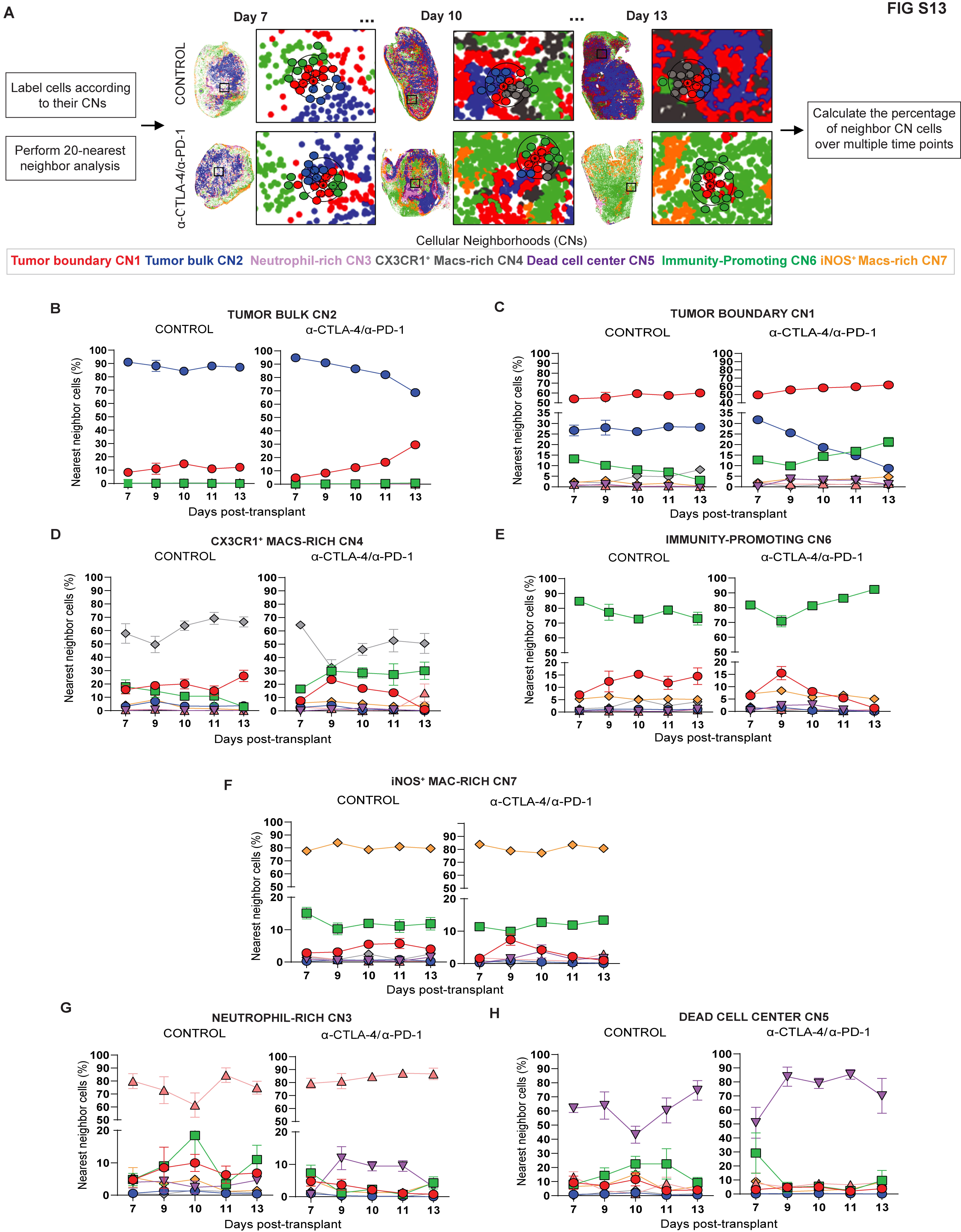

### Supplemental figure 14

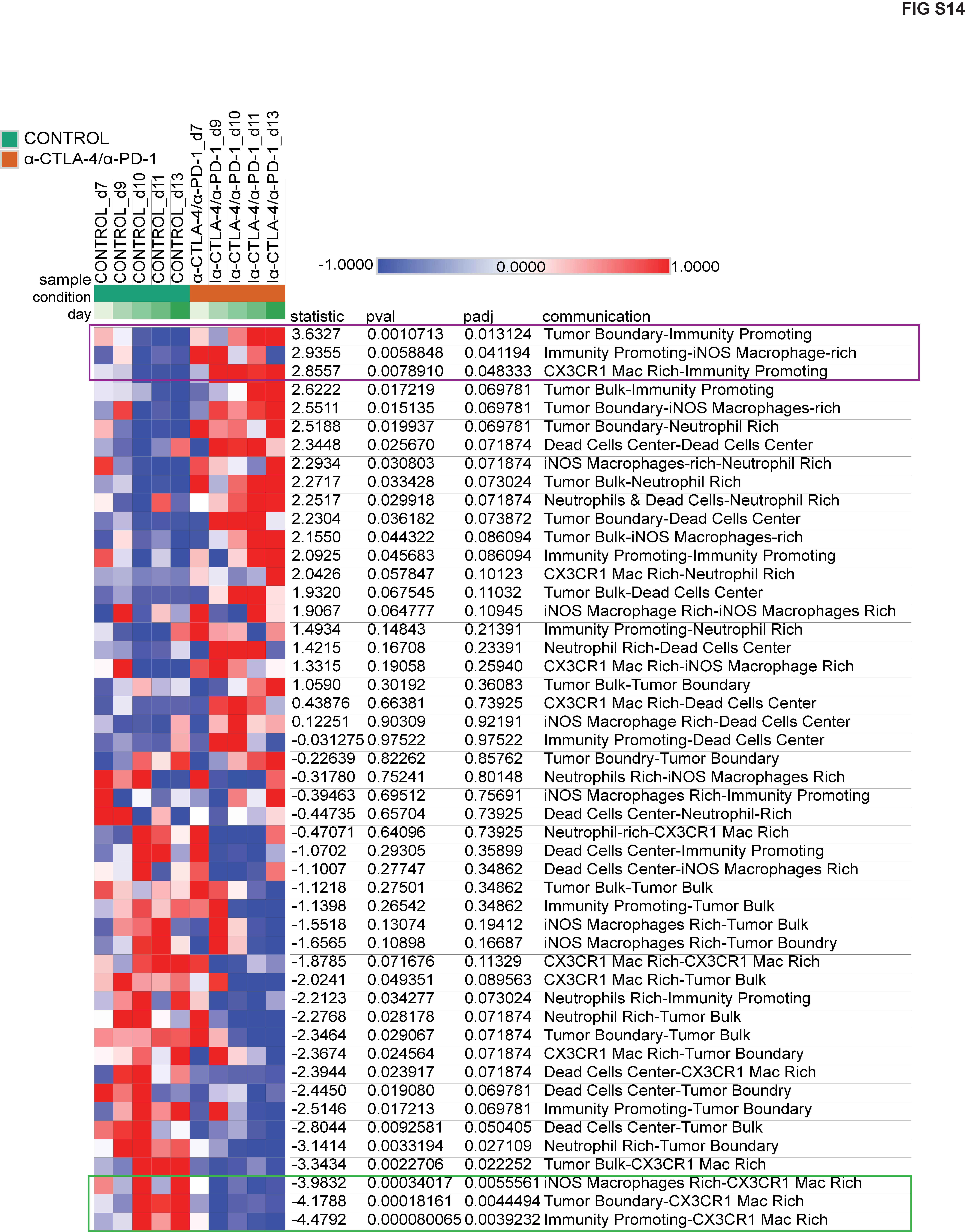
