## Supplemental Table 2 for "Immune Checkpoint Therapy Drives Maturation of a Cellular Neighborhood Nucleated by T Cell-APC Triads Enabling Spatially Compartmentalized Tumor Immunity"

| Cycle | Alexa 488 | ATTO 550 | Cy-5 |
|---|---|---|---|
| **1** | Blank | Blank | Blank |
| **2** | CD103-RX040 | ki67-RX047 | FOXP3-RX024 |
| **3** | CD140a-RX004 | CX3CR1-RX032 | CD3-RX021 |
| **4** | LY6C-RX037 | CD8a-RX029 | TBET-RX003 |
| **5** | NKP46-RX046 | TIM3-RX041 | XCR1 -RX045 |
| **6** | SIRP-alpha-RX013 | GrazymeB-RX017 | PD1-RX027 |
| **7** | CD206-RX031 | CD4-RX026 | CASPASE 3-RX015 |
| **8** | CD45-RX007 | CCR2-RX020 | Lag3-RX033 |
| **9** | Siglec F-RX016 | CD64-RX005 | F4/80-RX006 |
| **10** | CD38-RX019 | CD31-RX002 | CD11c-RX030 |
| **11** | CD24-RX022 | CD21-RX023 | INOS-RX036 |
| **12** | CD11b-RX025 | MHCII-RX014 | LY6G-RX042 |
| **13** | CD90.2-RX001 | empty | empty |
| **14** | PDL1-RX028 | empty | empty |
| **15** | Collagen 1-RX010 | empty | empty |
| **16** | ICOS-RX034 | empty | empty |
| **17** | Blank | Blank | Blank |

**Table S2. CODEX imaging cycles information.**
